# Dynamic proteomic profiling reveals protein-specific regulation of synthesis rates underpinning the divergent adaptation of human muscle to endurance and resistance training

**DOI:** 10.64898/2026.04.02.715579

**Authors:** Connor A. Stead, Aaron C.Q. Thomas, Priscilla Ma, Mackinley Szumlanski, Alysha C. D’Souza, James McKendry, Changhyun Lim, Jonathan C. McLeod, Chris McGlory, Ivo Siekmann, Jatin G. Burniston, Stuart M. Phillips

**Affiliations:** Research Institute for Sport & Exercise Sciences, Liverpool John Moore’s University, Liverpool. United Kingdom; Protein Metabolism Research Lab, Department of Kinesiology, McMaster University, Hamilton, ON. Canada; Food, Nutrition and Health, Faculty of Land and Food Systems, The University of British Columbia, Vancouver, BC, Canada; Edwin S.H. Leong Centre for Healthy Aging, Faculty of Medicine, The University of British Columbia, Vancouver, BC Canada; Population Health Sciences Institute, Faculty of Medical Sciences, Newcastle University, Newcastle upon Tyne. United Kingdom; School of Kinesiology and Health Studies, Queen’s Arts and Science, Queen’s University, Kingston, ON. Canada; School of Computer Science and Mathematics, Liverpool John Moores University, Liverpool, United Kingdom; Artificial Intelligence and Digital Technologies Research Institute (AIDTRI), Liverpool John Moores University, Liverpool, United Kingdom

**Keywords:** Protein turnover, Stable isotope labelling, Deuterium oxide, Skeletal muscle, Exercise

## Abstract

Endurance (END) or resistance exercise (RE) training results in adaptations that give rise to distinct skeletal muscle phenotypes. Hallmarks of RE include increases in muscle fibre and muscle cross-sectional area and strength, whereas END increases mitochondrial content. Such distinct phenotypes arise from differential metabolic and mechanical signal transduction, transcriptional, and protein translation pathways, culminating in exercise mode-specific adaptations in the muscle proteome. However, little empirical data exist on the protein-specific dynamic responses underlying training-mode-specific adaptations in humans. Using a model of unilateral exercise combined with stable isotope labelling with deuterium oxide, we measured changes in synthesis and abundance from baseline and during early (week 1) and later (week 10) periods of adaptation to END and RE training in young healthy adults (n = 14; 8 female, 6 male; 20 ± 1 y, 70 ± 10 kg). We quantified changes in the abundance (n = 1146 proteins) and synthesis (n = 247 proteins) profiles of skeletal muscle across a 5-day pre-training baseline period and during early and later adaptation to RE and END. Abundance profiling revealed mode-specific proteome remodelling, whereby RE increased ribosomal and contractile protein networks, whereas END increased mitochondrial inner membrane proteins after 10 weeks of training. The protein-specific synthesis rates of 119 proteins showed training-induced differences (P < 0.1 and log2 fold change > 1), including subsets of structural proteins that responded differently to RE and END training modes. Notably, distinct Z-disc proteins, such as XIRP1 (RE-specific) and LDB3 (END-specific), exhibited mode-specific regulation despite sharing a similar subcellular localisation. We report, for the first time, that divergent phenotypic adaptations to RE and END extend beyond changes in bulk fraction-specific synthesis rates and are regulated by training-mode-specific adaptations in distinct protein subsets within similar subcellular protein locations.

## Introduction

Skeletal muscle is a highly adaptable tissue capable of phenotypic alterations in response to changes in contractile activity. Resistance exercise (RE) results in muscle hypertrophy and improvements in muscle strength (Roberts et al., 2023, McLeod et al., 2024). In contrast, endurance exercise (END) leads to robust increases in mitochondrial content and capillary density within skeletal muscle, resulting in greater aerobic capacity (Mølmen et al., 2025, Bishop et al., 2019). A defining difference between RE and END, beyond differences in metabolic demand, is in the mechanical loading pattern imposed on the working muscle. END involves repetitive, low-load, submaximal contractions that generate relatively low forces across many cycles, with high metabolic demand. In contrast, RE imposes high-load, lower-frequency contractions that subject the sarcomere and its structural attachments to substantially greater peak mechanical stress. The differences in loading magnitude and pattern between END and RE are proposed to underpin the divergent adaptive outcomes of the two modes, yet the molecular pathways by which skeletal muscle transduces high mechanical load into the specific downstream signalling events that drive hypertrophic remodelling remain incompletely resolved (Wackerhage et al., 2019, Zhu et al., 2025). Mechanotransduction in skeletal muscle is likely to involve structural proteins at the sarcomere periphery, including the Z-disc, costameres, and extracellular matrix, given their position at the interface between the force-generating actin-myosin filaments and the intracellular signalling machinery (Burkholder, 2007)(Jorgenson et al., 2020). These structural compartments are therefore potentially load-sensing elements whose differential activation between training modes may encode information about the magnitude and pattern of mechanical input.

Muscle adaptation to repeated exercise is a fundamentally dynamic process, unfolding progressively over weeks of training rather than as a discrete event (Egan and Zierath, 2013). In humans, the timescale over which structural and metabolic remodelling becomes reliably detectable varies by training modality and outcome measure. Mitochondrial content, assessed by marker enzyme activity or volume density, increases detectably within 2–6 weeks of endurance training and continues to accumulate to a plateau with sustained training (Mølmen et al., 2025). Expansion of muscle fibre cross-sectional area in response to RE follows a slower trajectory (Damas et al., 2016), with detectable muscle fibre hypertrophy emerging only after approximately 10-12 sessions or ∼6-8wk of RE training (Goreham et al., 1999) and becoming robust after around 18 sessions of progressive loading (Damas et al., 2018). Understanding how this adaptation progression occurs and how it differs between END and RE training requires tracking molecular responses across different phases of a training intervention.

The divergent phenotypic outcomes of RE and END training are ultimately the product of protein-specific remodelling, in which the selective turnover of individual proteins reshapes the composition and functional capacity of the muscle proteome. Protein abundance profiling before and after exercise training consistently demonstrates substantial remodelling across metabolic, structural, and regulatory proteins in both humans and animal models (Hesketh et al., 2021, Emanuelsson et al., 2024), confirming that coordinated changes in proteome composition define adaptation. Importantly, untargeted discovery approaches have expanded the molecular landscape of adaptation beyond the canonical targets of myosin heavy chain isoforms and mitochondrial proteins, revealing that other proteome compartments, such as calcium-binding and cytoskeletal proteins, are also remodelled but were largely ignored in prior hypothesis-driven analyses (Hulmi et al., 2025). However, abundance profiling alone cannot resolve the dynamic processes driving remodelling. Steady-state protein abundance reflects the net balance of synthesis and degradation rather than providing information on how these processes contribute to the specificity of adaptation.

Dynamic proteome profiling combines stable isotope (deuterium oxide; D₂O) labelling in vivo with high-resolution peptide mass spectrometry and measures both the synthesis rates and abundances of individual skeletal muscle proteins across periods of days to weeks in free-living humans (Burniston, 2019). Dynamic proteome profiling reveals heterogeneity in protein turnover within subcellular compartments and can expose regulatory patterns that are invisible to bulk mixed-protein measurements (Stead et al., 2025b). To date, the majority of proteomic studies report either abundance data only /or protein-specific synthesis rates only. However, information on changes in protein synthesis rate without accompanying abundance data results in the inability to determine whether elevated synthetic rates reflect net accretion or accelerated renewal of an unchanged pool (i.e., protein turnover). For example, dynamic proteome profiling of protein-specific FSR and abundance revealed that increases in the synthesis of metabolic enzymes were not accompanied by increases in their abundance in response to high-intensity interval training (Srisawat et al., 2023), suggesting that training led to increased turnover rather than driving net accretion of these proteins. Studies highlight that the specificity of adaptation may be regulated at the level of individual proteins, rather than homogenous control of synthesis across all proteins comprising a bulk fraction (Camera et al., 2017, Stead et al., 2025a). Therefore, whether, and how, END and RE drive divergent muscle phenotypes through the selective synthesis of distinct proteins or through broadly uniform synthetic responses that differ only in magnitude remains unanswered.

In this study, we provide the first protein-by-protein characterisation of the dynamic proteome responses underlying divergent skeletal muscle adaptation to END and RE training in humans. Using a within-subject, unilateral design (MacInnis et al., 2017b), we quantified changes in protein synthesis and abundance across 3 phases: a 5-day free-living basal period, the first week (Early) and tenth week (Later) of END and RE training, enabling protein-specific FSR and abundances to be resolved across distinct phases of baseline, early, and later adaptation. We hypothesised that this approach would reveal divergent adaptations to END and RE, characterised by time-dependent, selectively regulated dynamics of individual proteins.

## Materials and Methods

### Ethical Approval

Before participation, participants provided written informed consent to the experimental procedures approved by the Hamilton Integrated Research Ethics Board (HIREB #15317). The study conformed to the standards for the use of human subjects in research as outlined by the Canadian Tri-Council Policy (TCPS 2 2022) on the ethical use of human subjects in research and the Declaration of Helsinki.

### Experimental Design

We used a within-subject repeated-measures design for unilateral exercise training; 14 participants (8 females, 6 males) were recruited. Participants were healthy, recreationally active, and aged 18 to 24. Anthropometric and physiological testing was completed >72 hours before the initiation of D_2_O labelling in the Baseline and Later study period. As well as in the place of a regular training session in week 5, to adjust exercise prescription if required. During each measurement period, participants were provided with a standardised weight-maintaining diet containing >1.4g protein/kg/d.

Participants’ legs were randomly assigned to undertake RE or END training. Biopsies from the vastus lateralis were collected at the start (unilateral sample collected on day 0) and end (bilateral samples collected on day 4) of the Baseline period. Participants refrained from moderate-vigorous exercise across the Baseline period. Immediately following the end of the Baseline period, participants began a 10-week unilateral exercise intervention that required training thrice weekly using unilateral lower-limb RE and END in contralateral limbs. Dynamic proteomic responses to training were quantified over the first 5 days (Early) and the final 5 days (Later) of RE and END, with bilateral biopsies collected at the start and end of the 1st and 10th weeks of training. Initial biopsies in each study period (Early and Later) were collected immediately before exercise, while biopsies at the end of each study period were collected 3 h after the final exercise bout. Biosynthetic labelling of proteins with D_2_O was via 2 days of loading doses and sustained consumptionq throughout the Baseline, Early, and Later study periods. Throughout the labelling periods, saliva samples were collected daily to measure precursor (whole-body water) enrichment.

### Physiological Assessment

Body composition and leg lean mass were assessed using dual-energy X-ray absorptiometry (DEXA) scanning. Subjects were fasted (∼10 h) and had not completed any exercise during the 48 h prior to the pre-training scan, prior to the initiation of the Baseline, Early, and Later measurement periods (Figure 1). Muscle thickness and CSA were assessed by ultrasonography following DEXA, as previously described (Stokes et al., 2021). In brief, participants lay supine for 10 minutes to normalise fluid shifts, with their feet positioned in a custom foot-hold apparatus that prevented their thighs from being depressed against the bed. Ultrasonography assessment occurred at a point equidistant between the greater trochanter of the femur and the lateral epicondyle of the knee, identified by palpation. Following ultrasound, participants performed unilateral (RE limb only) 1-repetition maximum strength tests of the quadriceps press and extension. Participants were instructed on proper lifting technique and performed a warm-up set of 10 repetitions at ∼50% 1RM. Participants provided feedback on the difficulty of each set using an adapted Borg Scale (CR-10) (Buckley and Borg, 2011), and the load was progressively increased by ∼ 5-20% for each repetition until a ‘true’ 1RM was achieved, as previously described (McGlory et al., 2014); participants were given 3 min rest between each attempt. Participants also completed a unilateral (END limb only) incremental exercise test to exhaustion on a cycle ergometer (Velotron, RacerMate; Seattle, WA) as previously described (Thomas et al., 2022). The test began with a 2 min warm-up at 50 watts (W), followed by incremental increases in resistance (1 W every 4 s) until volitional exhaustion or the point at which cycling cadence fell below 60 rpm. Maximal workload (W_max_) was recorded as the highest power output achieved during the last 30 s of the test.

**Figure 1.**
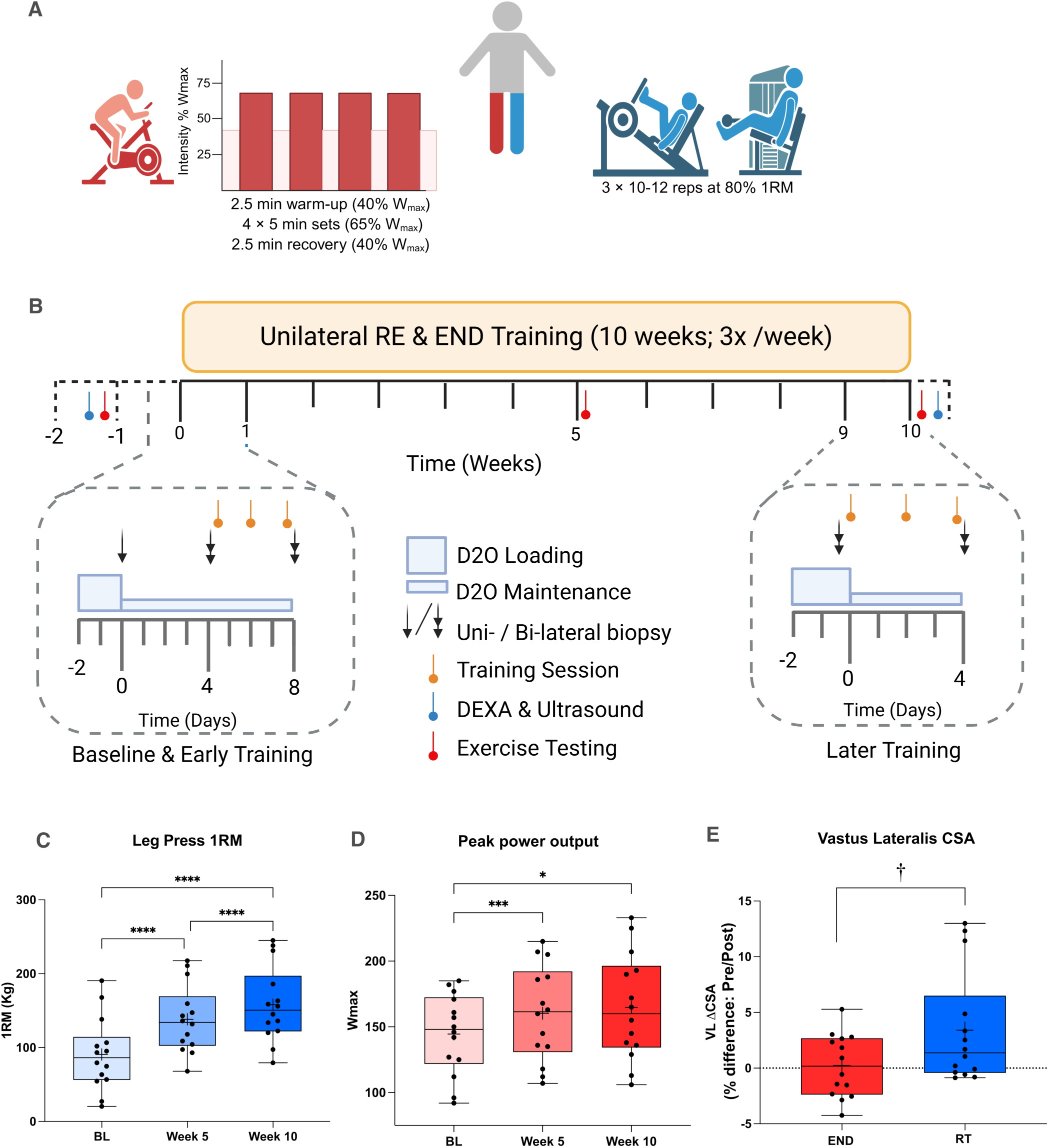
Experimental design. Experimental design of unilateral resistance and endurance training across 10-weeks. (A) Participants (n = 14; 8F/6M) were recruited, and legs were randomised to either resistance (RE) or endurance (END) exercise. (B) Participants then completed the 12-week study period, comprising dynamic proteome profiling across the baseline and early training periods, as well as the later training period. Participant physiological and exercise testing were measured at baseline, after 5 weeks and after 10 weeks for (C) leg press (1RM), (D) peak power output (watts) and (D) Ultrasound of the *vastus lateralis* measured at 50% the length of the femur. * (P<0.05), *** (P<0.001),**** (P<0.0001), † (P<0.01) Significantly different between groups.

### Unilateral Training Protocol

The training period consisted of participants visiting the lab 3 times per week for supervised exercise sessions. On each visit, subjects completed unilateral endurance and resistance training in a counterbalanced, randomised order. RE consisted of 3 sets × 10-12 reps at 80% 1RM leg press followed by 3 sets × 10-12 reps at 80% 1RM leg extension, with 2 min recovery between sets and exercises. END consisted of 2.5 min warm-up at 40% W_max_ followed by 4 bouts of 5 min cycling at 65% W_max_ with 2.5 min recovery (40% W_max_) between sets, followed by 2.5 min of ‘cooldown’ at 40% W_max_. Throughout each training session, participants were periodically monitored for heart rate (Polar A3, Lake Success, NY) and RPE (Borg CR10).

### Stable isotope labelling with deuterium oxide

We applied a two-phase approach to stable isotope labelling of newly synthesised proteins, consisting of a ‘loading phase’ and a ‘maintenance phase’ of D_2_O provision. Labelling was conducted on days -2 and -1 of each measurement period, during which participants consumed 10 doses of 0.66mL/kg body mass of 99.8 atom % deuterium oxide (Sigma-Aldrich) approximately 1 hour apart each day (equating to D_2_O loading of 13.2 mL/kg body mass). Precursor enrichment was then maintained by consumption of 1 daily dose of 0.66mL/kg body mass D_2_O during each measurement period. Body water enrichment was measured in saliva samples by cavity ring-down spectroscopy using a Liquid Water Isotope Analyser with an automated injection system (Los Gatos Research, Mountain View, CA). The water phase of the saliva was injected six times, and the average of the last three measurements was used for data analysis as described previously (McKendry et al., 2019).

### Muscle collection and processing

Muscle biopsies were collected as previously described (Wilkinson et al., 2014). Muscle samples (∼35 mg) were pulverised in liquid nitrogen, then homogenised on ice in 10 volumes of 1% Triton X-100, 50 mM Tris, pH 7.4, that contained complete protease inhibitor (Roche Diagnostics, Lewes, United Kingdom) using a PolyTron homogeniser. Samples were incubated on ice for 15 min, then centrifuged at 1000 *g*, 4°C, for 5 min. Supernatants that contained soluble/sarcoplasmic proteins were decanted and stored on ice, and the myofibrillar pellet was resuspended in 0.5 mL of homogenisation buffer, then centrifuged at 1000 *g*, 4°C, for 5min. The washed myofibrillar pellet was solubilised in 0.5 mL of 7 M urea, 2 M thiourea, 4% CHAPS, 30 mM Tris, pH 8.5, and cleared by centrifugation at 12,000 *g*, 4°C, for 45 min. Protein concentrations of both the myofibrillar fraction and sarcoplasmic fraction were measured by using the Bradford assay (Sigma-Aldrich, Poole, Dorset, UK). Aliquots that contained 100 μg protein were precipitated in 5 volumes of acetone for 1h at −20°C. Tryptic digestion was performed using the filter-aided sample preparation method (Wiśniewski et al., 2009). Aliquots containing 100 µg protein were precipitated in acetone and resuspended in 40 μl UA buffer (8 M urea, 100 mM Tris, pH 8.5). Samples were transferred to filter tubes and washed with 200 µl of UA buffer. Proteins were incubated at 37 °C for 15 min in UA buffer containing 100 mM dithiothreitol, followed by incubation (20 min at 4 °C) protected from light in UA buffer containing 50 mM iodoacetamide. The UA buffer was exchanged with 50 mM ammonium bicarbonate, and sequencing-grade trypsin (Promega, Madison, WI, USA) was added at an enzyme-to-protein ratio of 1:50. Digestion was allowed to proceed at 37 °C overnight, and the peptides were collected in 100 μl of 50 mM ammonium bicarbonate containing 0.2 % trifluoroacetic acid. Samples containing 4 µg of peptides were desalted using C18 Zip-tips (Millipore) and resuspended in 20 µl of 2.5 % (v/v) acetonitrile (ACN), 0.1 % (v/v) formic acid (FA) containing 10 fmol/μl yeast alcohol dehydrogenase (ADH1; MassPrep, Waters Corp., Milford, MA).

### Liquid chromatography tandem-mass spectrometry

Peptide mixtures were analysed using an Ultimate 3000 RSLC nano liquid chromatography system (Thermo Scientific) coupled to a Q-Exactive Orbitrap mass spectrometer (Thermo Scientific). Samples were loaded onto the trapping column (Thermo Scientific, PepMapTM 100, 5 μm C18, 300 μm X 5 mm), using ulPickUp injection, for 1 minute at a flow rate of 25 μl/min with 0.1 % (v/v) TFA and 2% (v/v) ACN. Samples were resolved on a 110 cm analytical column (µPAC Column; Thermo Fisher) using a gradient of 97.5 % A (0.1 % formic acid), 2.5 % B (79.9 % ACN, 20 % water, 0.1 % formic acid) to 50 % A 50 % B over 90 min at a flow rate of 300 nl/min. The data-dependent selection of the top 10 precursors over the m/z range 300-1600 was used for data acquisition, consisting of a 70,000-resolution full-scan MS scan at m/z 200 (AGC set to 3e^6^ ions, maximum fill time 240 ms). MS/MS data were acquired using quadrupole ion selection with a 3.0 m/z window, HCD fragmentation with a normalised collision energy of 30 and in the Orbitrap analyser at 17,500-resolution at m/z 200 (AGC target 5e^4^ ions with a maximum fill time of 80 ms). To avoid repeated selection of peptides for MS/MS, the program used a 30 s dynamic exclusion window.

### Dynamic Proteome Profiling

Progenesis Quantitative Informatics for Proteomics (QI-P; Nonlinear Dynamics, Waters Corp., Newcastle, UK) was used for label-free quantitation, consistent with previous studies (Camera et al., 2017, Hesketh et al., 2020, Burniston et al., 2014). Log-transformed MS data were normalised by inter-sample abundance ratio, and relative protein abundances were calculated using non-conflicting peptides only. In addition, abundance data were normalised to the 3 most abundant peptides of yeast ADH1 to obtain abundance estimates in fmol/ µg protein. MS/MS spectra were exported in Mascot generic format and searched against the Swiss-Prot database (2022_08) restricted to ‘Homo Sapiens (20,371 sequences) using a locally implemented Mascot server (v.2.8 www.matrixscience.com). The enzyme specificity was trypsin with 2 allowed missed cleavages, carbamidomethylation of cysteine (fixed modification), deamidation of asparagine and glutamine (variable modification) and oxidation of methionine (variable modification). M/Z error tolerances of 10 ppm for peptide ions and 0.6 Da for fragment ion spectra were used. The Mascot output (XML format), restricted to non-homologous protein identifications, was recombined with MS profile data in Progenesis. Protein abundance data were collected for unique peptides with an identification score < 1% false discovery rate.

Mass isotopomer abundance data were extracted from MS spectra using Progenesis Quantitative Informatics (Nonlinear Dynamics, Newcastle, UK). Consistent with previous work (Camera et al., 2017, Hesketh et al., 2020, Nishimura et al., 2023), the abundances of peptide mass isotopomers were collected over the entire chromatographic peak for each proteotypic peptide that was used for label-free quantitation of protein abundances. Mass isotopomer information was processed using in-house scripts written in Python (version 3.12.4). The incorporation of deuterium into newly synthesised protein was assessed by measuring the increase in the relative isotopomer abundance (RIA) of the m1 mass isotopomer relative to the sum of the m0 and m1 mass isotopomers (Equation 1) that exhibits rise-to-plateau kinetics of an exponential regression (Sadygov, 2020) as a consequence of biosynthetic labelling of proteins in vivo.

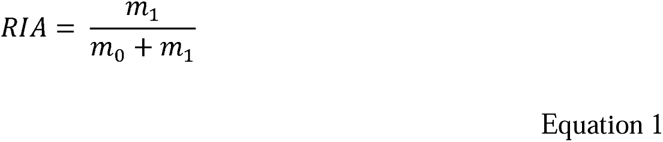

The plateau in RIA (RIAplateau) of each peptide was derived (Equation 2) from the total number (N) of 2H exchangeable H—C bonds in each peptide, which was referenced from standard tables (Holmes et al., 2015), and the difference in the D:H ratio (2H/1H) between the natural environment (DHnat) and the experimental environment (DHexp) based on the molar percent enrichment of deuterium in the precursor pool, according to (Ilchenko et al., 2019).

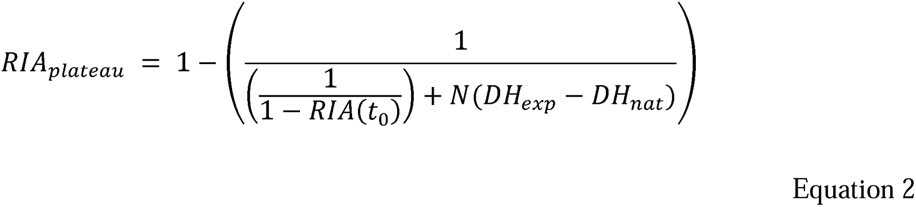

The rate constant of protein degradation (kdeg) was calculated (Equation 3) between the beginning (t0) and end (t1) of each 10-day labelling period. Calculations for exponential regression (rise-to-plateau) kinetics reported previously (Ilchenko et al., 2019) were used, and kdeg data were adjusted for differences in protein abundance (P) between the beginning (t0) and end (t1) of each labelling period.

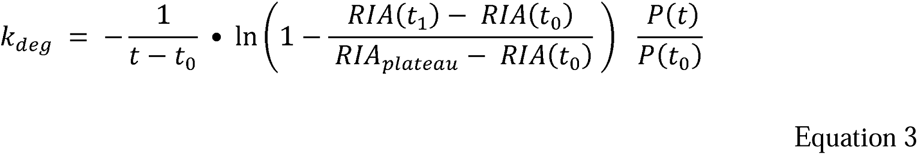

### Bulk Fractional Protein Synthesis rates

Muscle samples (∼30–50 mg) were homogenised on ice in buffer [10 μL/mg 25 mM Tris-HCl pH7.2 0.5% vol:vol Triton X-100 and protease/phosphatase inhibitor cocktail tablets (Complete Protease inhibitor Mini-Tabs, Roche; and PhosSTOP, Roche Applied Science)] and centrifuged at 2250g for 10 min at 4°C to separate the sarcoplasmic and myofibrillar enriched protein fractions. The supernatant was removed for sarcoplasmic protein analysis, and the myofibrillar pellet was retained (McKendry et al., 2024). Doubly distilled water (500 μL) was added to the myofibrillar pellet, which was vortexed and centrifuged (250g for 10 min at 4°C). The supernatant was removed and discarded; the remaining pellet was incubated in 0.3 M NaOH for 30 min at 50°C to solubilise the myofibrillar protein fraction. The samples were vortexed and centrifuged at 11200g for 5 min at 4°C, supernatant (myofibrillar protein) was removed, and the collagen pellet was discarded. This step was repeated, and supernatants were pooled. Myofibrillar and sarcoplasmic proteins were precipitated with 1 mL of 1 M perchloric acid and centrifuged at 700g for 10 min at 4°C. The supernatant was discarded, and the fraction was washed twice with 70% ethanol. Ethanol was removed, and the sample was placed in 1mL of 1 M activated Dowex H+ (50XW8 Cation exchange resin) and 1mL of 1 M HCl. The protein–enriched pellets were hydrolysed at 110 °C for 72 h to release their respective amino acids (AAs). Following hydrolysis, the AAs were purified by ion-exchange chromatography on Dowex H+ resin using 2 M NH_4_OH, and the N (O,S)-ethoxycarbonyl ethyl ester of alanine was prepared as described previously (Husek, 1991). 2H-alanine enrichment was measured by Trace 1310 Gas Chromatrography – GC Isolink II Pyrloysis - Delta Q Isotope Ratio Mass Spectrometry (Thermo Scientific). The data were normalised to the VSMOW scale (‰ vs VSMOW) using an alkane mix with known certified deuterium enrichments (Indiana Mix B5, © Schimmelmann Standards LLC), and internal standards of D4-Alanine enrichment powders were run alongside samples to account for machine drift. 1uL of sample was injected in splitless mode four times, and the best three injections were used for downstream analyses. The average coefficient of variation for each sample across three injections was 0.5% (range: 0.0% - 7.7%). Myofibrillar protein synthesis (MPS) was calculated using the standard precursor-product equation (Equation 4).

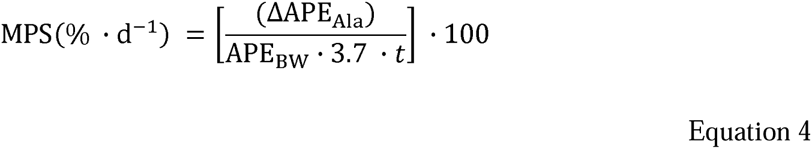

Where D APE_Ala_ is the change in protein-bound 2H-alanine enrichment between biopsy sampling points, APE_BW_ is the body-water 2 H enrichment multiplied by 3.7 to correct for the number of carbon-hydrogen bonds labelled in alanine relative to total body water, and t is the incorporation time (in days) between biopsies. Exercise-induced changes in MPS were calculated using the muscle deuterium enrichment values obtained from the muscle specimen taken from the same limb at Baseline, Early and Later weeks of training.

### Statistical and Bioinformatic Analysis

A one-way analysis of variance was applied to investigate the physiological responses to training at Baseline, week 5 and week 10 of the training period (1RM and Watt peak testing). A t-test was used to investigate differences in physiological data (e.g., leg lean mass and muscle cross-sectional area) across exercise modes. Where appropriate, significant (p < 0.05) differences in physiological data between means were analysed using Tukey’s post-hoc analysis. A two-way analysis of variance was applied to investigate the Mixed MPS response to training across the Baseline, Week 1 and Week 10 periods. Where appropriate, significant (p < 0.05) differences in physiological data between means were analysed using Tukey’s post-hoc analysis.

Protein abundance data were stringently filtered to include proteins quantified in all sampling time periods (Baseline, Early and Later), timepoints (Pre and Post) and conditions (END and RE) in all n = 14 individuals. For dynamic proteomic data, to leverage the strength of within-subject comparisons of dynamic responses to different training modes over time, data were included only if quantified across all conditions (BL, Early END, Early RE, Later END, and Later RE) in≥ 4 participants. Linear modelling of protein abundance and synthesis data was conducted using the LIMMA (Linear Models for Microarray Data) package in R. To account for repeated measures within participants, the duplicateCorrelation() function was used to estimate intra-participant correlations, which were incorporated into the model. All data were log₂-transformed prior to analysis to stabilise variance across the dynamic range. Empirical Bayes moderation was applied using eBayes() with both robust = true and trend = true, to enhance variance estimation. To identify changes in abundance and synthesis rate between exercise and baseline measurements, predefined contrasts were constructed to compare each post-training condition with baseline. In dynamic proteomic analyses, Type I error was assessed using false-discovery rate (FDR) estimates (q values) and BH-corrected P values using the ‘qvalue’ package (R/Bioconductor) (Storey and Tibshirani, 2003).

To aid in confirming divergence in muscle adaptation to RE and END, significant changes in protein abundance were considered when P < 0.05. Proteins showing significant differences in abundance were analysed for functional enrichment using the clusterProfiler package in R (Yu et al., 2012). UniProt accessions were mapped to Entrez IDs, with all quantified proteins used as the background universe. Over-representation analysis of Gene Ontology (GO) Cellular Component terms was performed using enrichGO. Enriched categories were visualised using enrichplot, and relationships between genes and enriched terms were displayed with cnetplot, showing the top 10 categories. Node colour represents protein-specific log₂ fold-change values.

Hierarchical clustering of dynamic proteomics data was performed using Ward’s minimum-variance method and Euclidean distance. To enable data-driven discovery of biologically meaningful patterns, proteins exhibiting changes in protein-specific synthesis with P < 0.1 and log2 fold-change > 1 were selected for further analysis. Groups of proteins meeting the criteria for further bioinformatic analysis were investigated using bibliometric mining in the Search Tool for the Retrieval of INteracting Genes/proteins (STRING, Version 12) using the evidence of interaction sources of bibliometric text mining, experimental verified protein-protein interaction data, gene ontology databases, and co-expression data with the minimum required interaction score set at 0.4 (medium confidence). Protein-protein interaction networks and functional enrichment analyses were conducted using the STRINGdb package in R (version 2.12.1) (Szklarczyk et al., 2019) and corrected for the experiment-specific background, comprising all proteins included in the statistical analysis. Protein-protein interaction networks were transferred using the RCy4 R package and visualised in Cytoscape version 3.10.3 (Shannon et al., 2003).

## Results

### Physiological response to unilateral training

Fourteen (8F/6M) young (20 ± 1 years), healthy, recreationally active individuals were recruited and completed a 10-week (3 sessions per week) unilateral END and RE protocol (Figure 1A). Participants’ physiological data are reported in Table 1. All participants completed ≥90% (27 of 30) of the programmed training sessions. Ten weeks of unilateral RE significantly increased leg press 1RM (p < 0.0001; mean diff = 67 ± 3 kg; Figure 1B) and knee extension 1RM (p <0.0001; mean diff = 13 ± 2 kg). END significantly (p = 0.035) increased unilateral W_max_ (20 ± 7 W; Figure 1C). Ten weeks of RE training resulted in a significant (p = 0.0002) increase in DEXA-measured leg lean mass in the RE limb only. Ultrasound imaging of the *vastus lateralis* also demonstrated a modest yet significant 12.6% increase (p = 0.01, mean diff = 3.4 cm^2^) in CSA in the RE limb only following 10 weeks of training (Figure 1D).

**Table 1.**
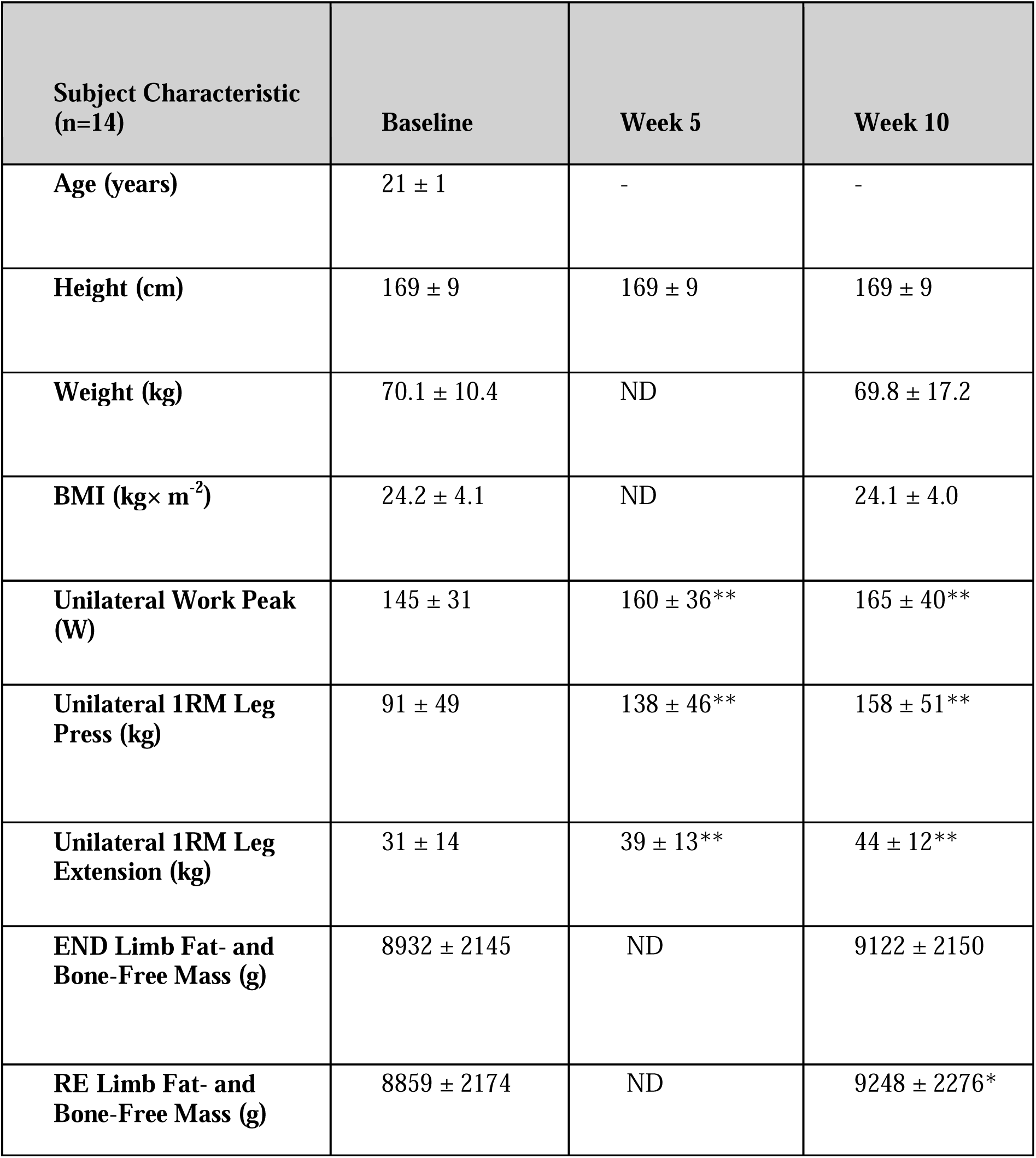
Physical and physiological characteristics were measured at baseline, after 5 weeks and 10 weeks (Post). Values are means±SD. Data shown for Work Peak are only from the END leg, and data for 1RM are only for the RE leg. Data are presented as means ± standard deviation. *p<0.05; **p<0.005 from Baseline. ND – not determined.

### Mixed-protein synthesis responses to unilateral training

Figure 2A shows the mixed (all protein) muscle protein synthesis responses. Both END and RE increased muscle protein synthesis responses in Early, which were still maintained in Later, but only in the RE leg.

**Figure 2.**
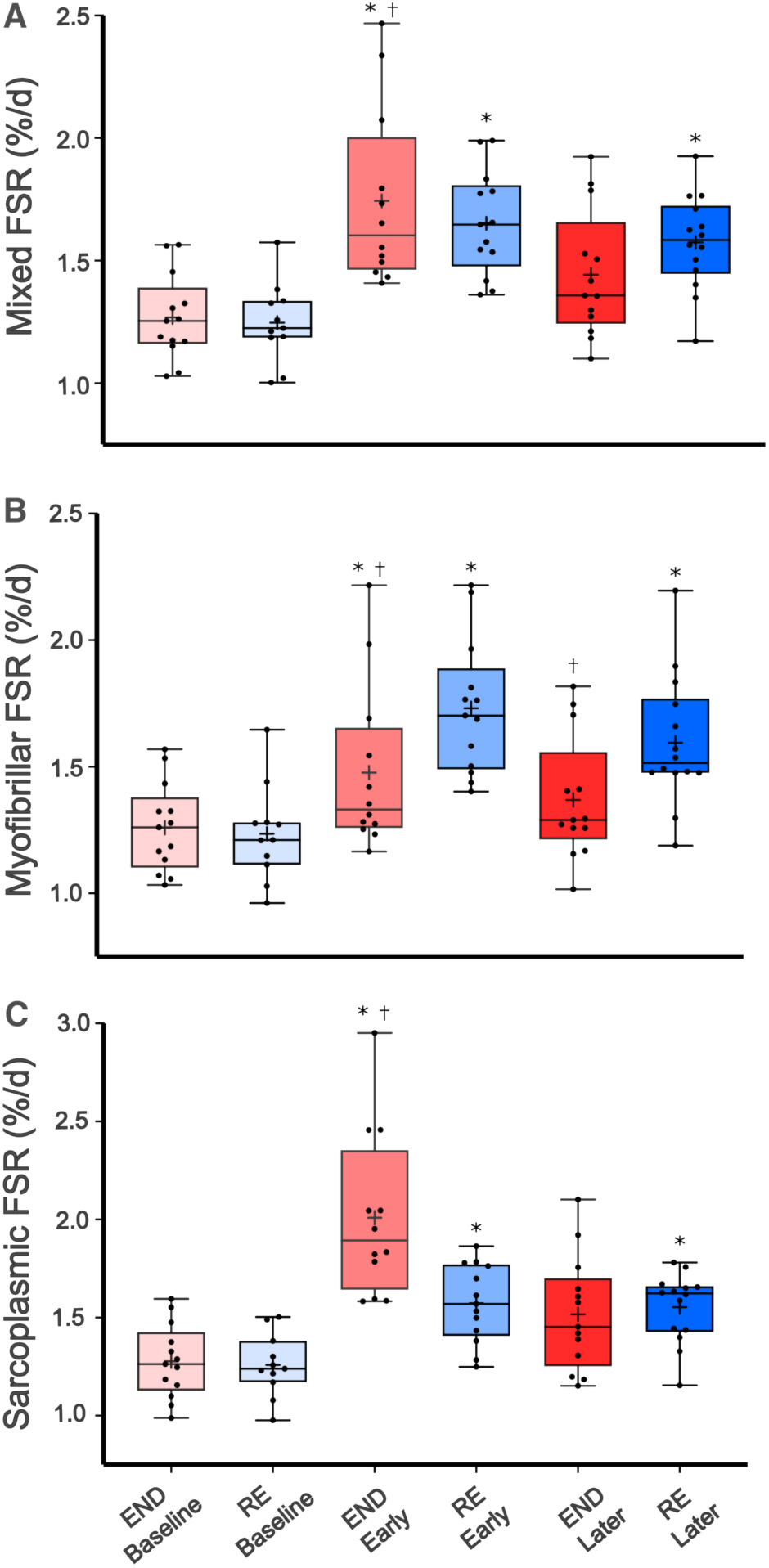
Changes in mixed protein fractional synthesis rate in response to unilateral exercise training. (A) Mixed (Bulk), (B) Myofibrillar and (C) Sarcoplasmic protein FSR, expressed as %/d, for the baseline (BL), 1st and 10^th^ week of exercise. † (P<0.01) Significantly different from RE at the same time point. * (P<0.01) Significantly different from baseline.

As shown in Figure 2B, myofibrillar protein synthesis increased during early END (1.48 ± 0.31 % day^-1^, p<0.01) and RE (1.73 ± 0.26 % day^-1^, p<0.0001) compared to baseline (1.24 ± 0.18 % day^-1^). Chronic RE resulted in sustained increases in myofibrillar protein synthesis during later adaptation (1.60 ± 0.26 % day^-1^, p < 0.001), whereas END did not differ compared to baseline (1.24 ± 0.18 % day^-1^).

Sarcoplasmic protein synthesis (Figure 2C) was increased in both early END (2.01 ± 0.40 % day^-1^, p<0.0001) and RE (1.57 ± 0.19 % day^-1^, p=0.0071) compared to Baseline (1.26 ± 0.15 % day^-1^). In the RE-trained limb, there was a significant increase in sarcoplasmic protein synthesis during week 10 (1.55 ± 0.17 % day-1) compared with Baseline (p<0.01). In the END limb, sarcoplasmic protein synthesis during week 10 (1.52 ± 0.29 % day^-1^) was not significantly different from Baseline.

### Changes in protein abundance indicate divergent adaptation to endurance and resistance training

Overall, 16,590 unique peptides comprising 1849 protein abundances were quantified across our entire protein abundance profiling (n ≥ 1 sample). To ensure robustness of our data, protein abundance data were stringently filtered to include proteins quantified in all 9 samples, encompassing all time periods (Baseline, Early, and Later), timepoints, and conditions (END and RE) for each of the n = 14 individuals. Altogether, protein abundance was determined for 1145 proteins quantified across all (n = 126) muscle samples.

Statistical analysis of changes in protein abundance after Early or Later END and RE revealed 254 significant changes (P < 0.05). To resolve global patterns in changes in protein abundance across time following RE and END, we performed enrichment analysis of significantly up- and downregulated proteins (Figure 3).

**Figure 3.**
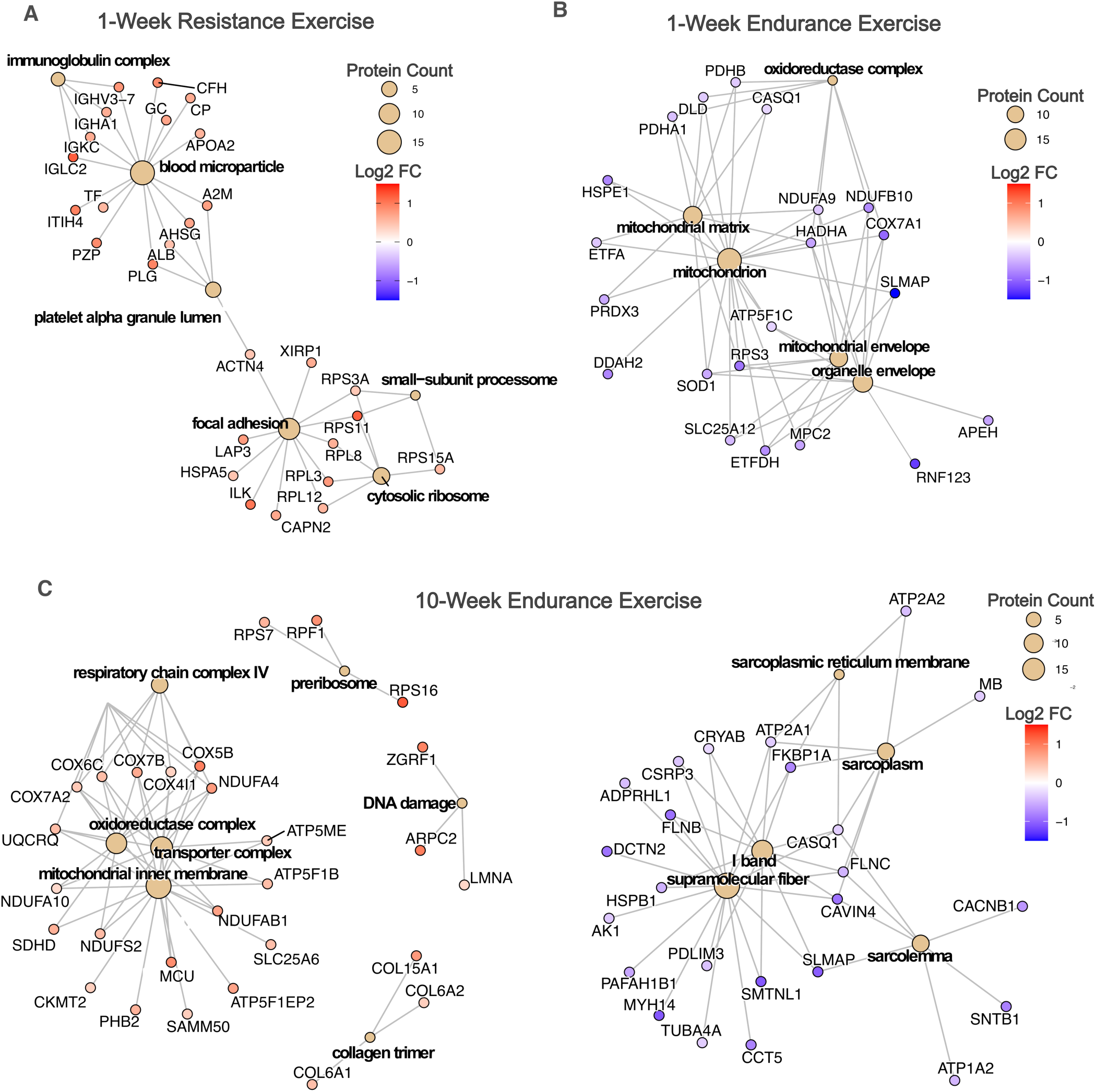
Changes in protein abundance resolve divergent adaptation to unilateral endurance and resistance training. Network plots of enriched Gene Ontology (GO) cellular component (CC) terms for proteins altered (P < 0.05) in abundance following resistance exercise (RE) and endurance exercise (END). (A) 1-week RE, (B) 1-week END, (C) 10-week END. No significant enrichments were detected after 10-week RE. Networks were generated from significantly enriched GO CC terms (FDR < 0.2). In each panel, the central yellow nodes with bold labels represent enriched GO terms. Peripheral connected nodes labelled with gene symbols represent proteins annotated to that term and detected as significantly different following training. Protein nodes are coloured by log₂ fold-change in abundance relative to baseline. GO term node size reflects the number of proteins annotated to that category. These networks highlight mode- and time-dependent differences in proteome organisation, including early ribosomal and adhesion-related enrichment in RE (A), early mitochondrial matrix and envelope remodelling in END (B), and later enrichment of respiratory chain, collagen, and cytoskeletal fibre networks in END (C).

Forty-six proteins were increased in abundance following 1-week of RE. Network analysis revealed enrichment of GO terms, revealing increased abundance of proteins associated with the cellular component ‘blood microparticle’ (15 proteins; FDR < 0.01). Also, networks of proteins associated with the ribosome (6 proteins; FDR = 0.11) and focal adhesion pathways (11 proteins; FDR = 0.07) were increased in abundance after 1-week of resistance training. No enriched protein networks were identified among proteins decreased in abundance after 1 week of RE (Figure 3A). Similarly, although 74 proteins were altered after 10 weeks of RE (34 increased and 40 decreased), network analysis revealed no significant enrichment of coordinated protein pathways at this later stage of adaptation to RE.

Unexpectedly, 1 week of endurance training resulted in decreased abundance of 34 proteins. Downregulated protein abundances after early END were enriched for proteins associated with the cellular component “Mitochondrion” (19 proteins; FDR = 0.07) with the majority (12/19) of these mitochondrial proteins decreased in abundance associated with the “Mitochondrial matrix” (FDR = 0.013; Figure 3B). No protein networks were enriched (FDR ≥ 0.5) among the 16 proteins increased in abundance following 1-week END. As expected, 10 weeks of END training resulted in increases in the abundance of 45 proteins, which were significantly enriched for GO cellular component ‘mitochondrion’ (25 proteins; FDR = 0.007), which was predominantly comprised of proteins associated with the ‘inner mitochondrial membrane’ (19 proteins; FDR < 0.001) (Figure 3C). Furthermore, 10-weeks END resulted in increased abundance of collagen proteins, including Collagen alpha-1(XV) chain, Collagen alpha-2(VI) chain, Collagen alpha-1(VI) chain, and Collagen alpha-1(IV) chain. In contrast, proteins that decreased after 10-weeks of END (53 proteins) were enriched for GO terms associated with muscle structure and cytoskeleton organisation, including ‘I Band’ (12 proteins, FDR = 0.008) and ‘supramolecular fibre’ (19 proteins, FDR = 0.0278) (Figure 3C). Complete statistical results for all proteins quantified and compared between baseline, early (1 week), and chronic (10 week) training responses are provided in Supplementary Table 2, with volcano plots illustrating the magnitude and significance of changes visualised in Supplementary Figure.

### Dynamic protein responses to exercise training

While abundance profiling captures the changes in proteome state before and after training, protein synthesis measurements provide mechanistic insight into the dynamic processes driving those changes. Importantly, these datasets differ because abundance is measured independently at pre- and post-training timepoints, allowing comparisons of proteome snapshots across time. Whereas fractional synthesis rates are calculated from D₂O incorporation between the pre- and post-training biopsies, capturing the rate of newly synthesised proteins measured during the training period. Together, these complementary approaches reveal both the outcome of adaptation (changes in abundance) and the processes governing that remodelling (synthesis rates).

Deuterium oxide ingestion resulted in stable body water enrichment, averaging 1.88 ± 0.15% during the Baseline and Early period and 1.85 ± 0.12% during the Later measurement period. In total, dynamic proteome profiling quantified protein-specific fractional synthesis rates (FSR) for 1556 individual proteins. To maximise power within the unilateral design, only proteins consistently quantified across baseline, Week 1 and Week 10 of both END and RE in ≥ 4 individuals were retained. This stringent quality-control filter reduced the protein list to 247 proteins, but it is necessary for correct within-subject statistical and bioinformatic analyses.

GO and KEGG (Kyoto Encyclopedia of Genes and Genomes) enrichment analysis of proteome coverage from our 247 proteins used in our analysis revealed strong representation of terms associated with fundamental muscle structure and function, including contractile fibre, myofibril, and Z-disc, alongside mitochondrial and metabolic pathways such as oxidative phosphorylation, aerobic respiration, and ATP metabolic processes. Therefore, confirming that our dynamic proteomic dataset encompasses key components of muscle biology and exercise adaptation.

At baseline, protein-specific synthesis rates spanned from 25.43 ± 32.76 %·d⁻¹ for Nidogen-1 (NID1) to 0.86 ± 0.65 %·d⁻¹ for myoglobin (MYG) (Figure 4A). Among 12 myosin-related, muscle-specific proteins detected, considerable heterogeneity in synthesis rates was observed, ranging from 1.63 %·d⁻¹ for myosin regulatory light chain 2 (MLRS) to 8.26 %·d⁻¹ for myosin-binding protein H (MYBPH) (Figure 4A). These findings highlight substantial variation in protein turnover within human skeletal muscle, both across the proteome and within functionally related protein subcompartments.

**Figure 4.**
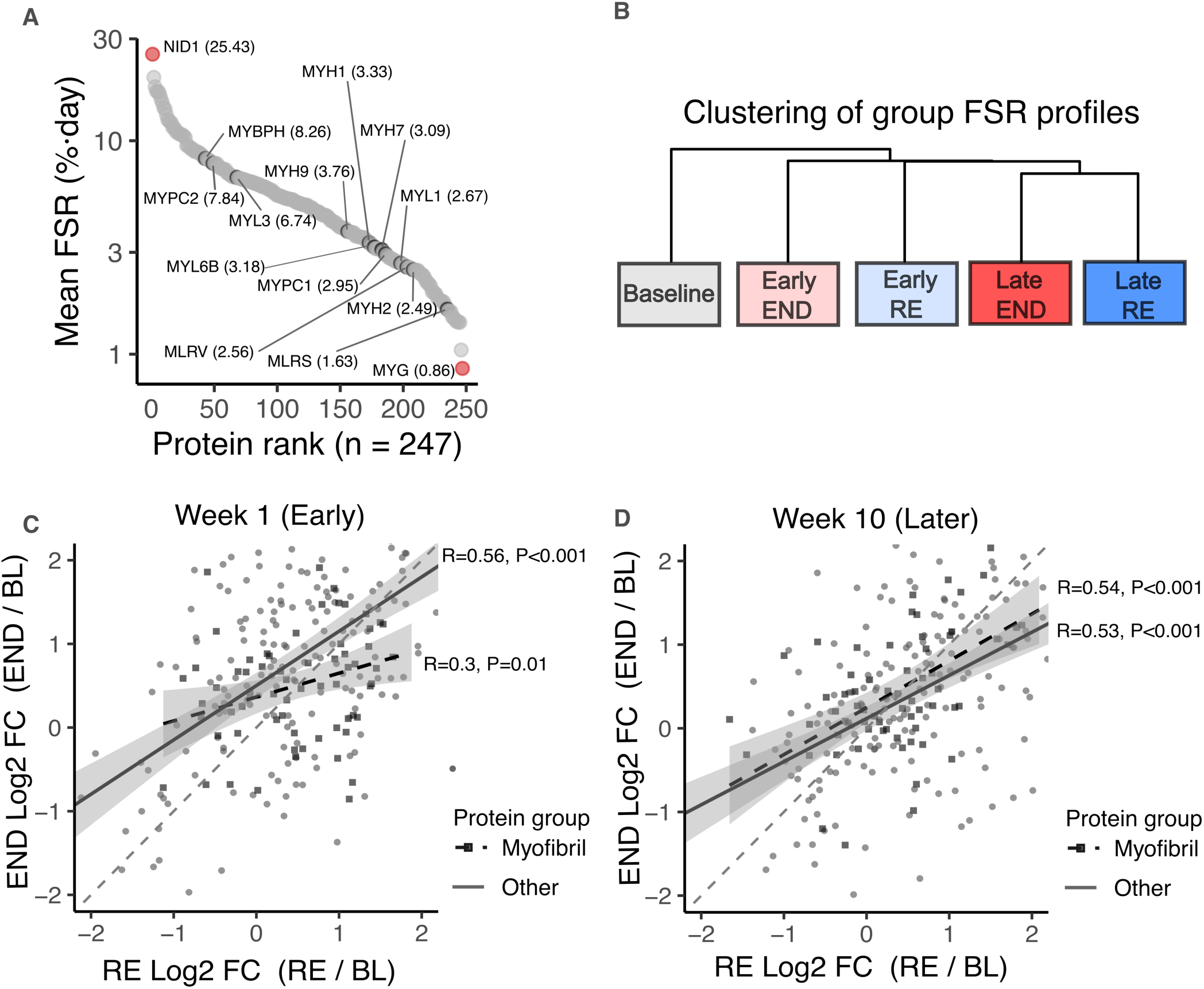
Dynamic proteome profiling of human skeletal muscle responses to endurance and resistance training. (A) Distribution of mean protein fractional synthesis rates (FSR; % · day⁻¹) across 247 quantified proteins ranked by mean rate. Key myosin-associated myofibrillar proteins (e.g. MYBPH, MYH isoforms, MYL, MYPC) are highlighted to illustrate the wide range of protein-specific turnover rates amongst similar protein functional classes. (B) Hierarchical clustering of group FSR profiles shows separation of early (Week 1) and late (Week 10) training phases, with resistance (RE, blue) and endurance (END, red) conditions diverging from baseline (grey). (C–D) Relationships between protein-specific synthesis responses (log₂ fold-change from baseline) for RE and END at Week 1 (C, Early) and Week 10 (D, Later). Each point represents an individual protein; myofibrillar proteins are indicated in black squares, all other proteins in grey circles. Solid and dashed lines denote regression lines for all proteins and myofibrillar proteins, respectively. Early adaptation showed a weaker correlation between modes (R = 0.30, P = 0.01) for myofibrillar proteins compared with later adaptation (R = 0.53 and 0.54, P < 0.001), indicating that while both modes increase overall synthesis rates, the specific subsets of proteins remodelled differ early in training before seemingly converging with continued adaptation.

Unsupervised hierarchical clustering of dynamic proteome profiles revealed that conditions predominantly grouped by training timepoint rather than exercise mode, with Week 1 END and RE clustering together and Week 10 END and RE clustering together (Figure 4B). Direct comparison of protein-specific synthesis responses between training modes revealed moderate overall correlations in mode-specific responses across the Early and Later periods. Across the proteome, fractional synthesis rates during Early adaptation were moderately correlated between RE and END (R = 0.5, P ≤ 0.0001), indicating that both modes stimulated broad increases in protein synthesis but not uniformly across all proteins. This relationship weakened when restricted to myofibrillar proteins (R = 0.3, P = 0.01), suggesting mode-specific differences in synthesis rates of distinct myofibrillar subsets during the first week of training (Figure 4C). In contrast, correlations between RE and END at Later adaptation were of similar magnitude (R = ∼ 0.5 for both myofibrillar and other proteins) (Figure 4D).

Differential analysis using LIMMA identified proteins with altered synthesis relative to baseline within each training mode. To focus on biologically meaningful changes, proteins were retained if they met thresholds of P < 0.1 and log₂ fold-change > 1 (equating to a 2-fold increase or decrease in synthesis rate). Volcano plots revealed broad proteome-wide responses during the early training period, whereas fewer proteins were significantly altered at Week 10 (Figure 5-6).

**Figure 5.**
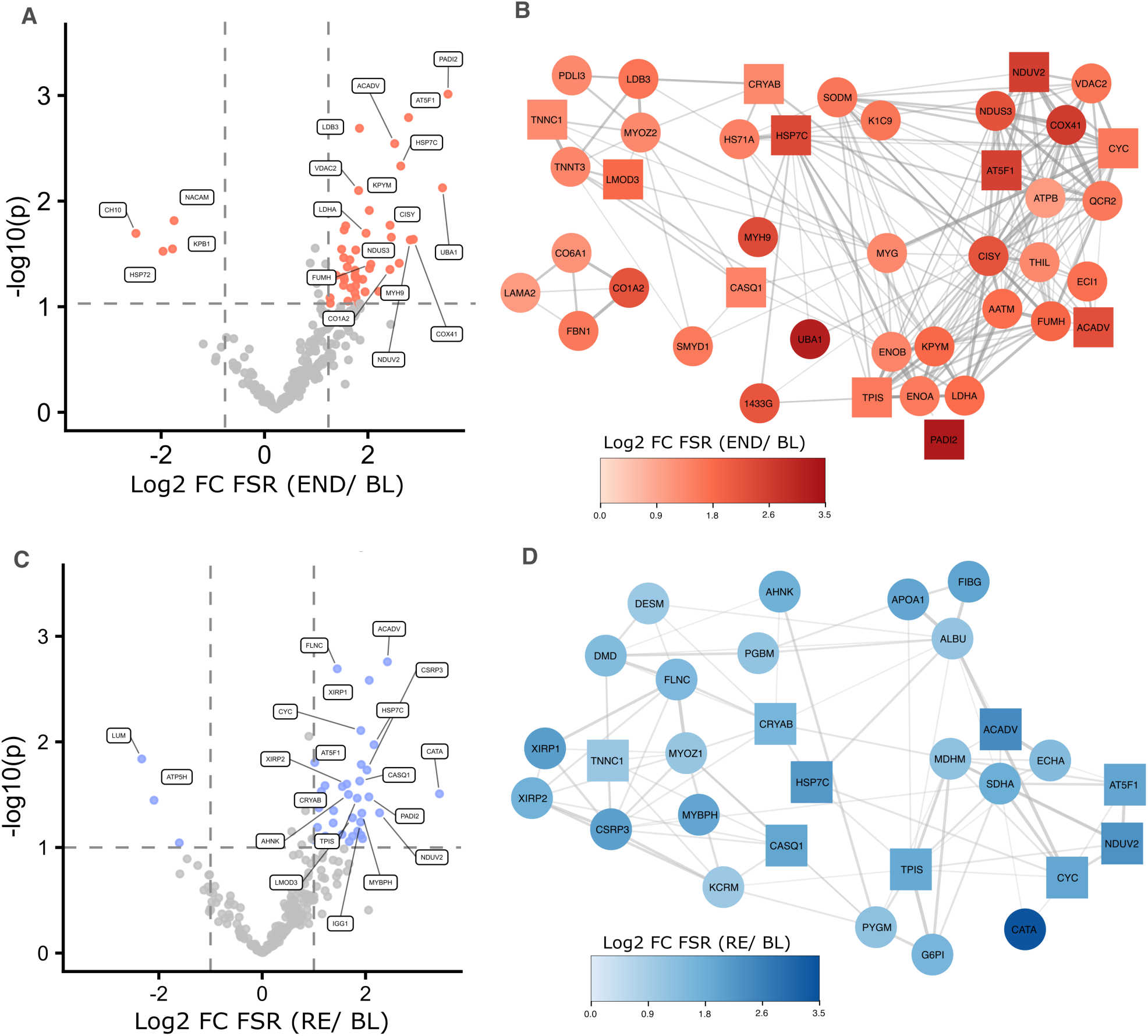
Mode-specific and shared upregulated proteins after one week of endurance (END) and resistance (RE) training. (A, C) Volcano plots showing protein-specific changes in fractional synthesis rate (FSR; log₂ fold-change vs baseline) following END (A) and RE (C). Significantly regulated proteins (p < 0.1 and log₂FC > 1) are coloured red (END) or blue (RE); grey points indicate non-significant proteins. Labelled proteins represent the top 20 significantly changed FSRs (log₂FC and –log₁₀ p). Dashed lines mark significance thresholds. (B, D) STRING protein–protein interaction networks of upregulated proteins after END (B) and RE (D). Node colour intensity reflects log₂ fold-change in FSR relative to baseline. Circles represent proteins uniquely upregulated in each mode, while squares indicate proteins common to both END and RE. Networks illustrate mode-specific clustering of cytoskeletal, metabolic, and chaperone proteins, with limited overlap in the specific subsets remodelled by each training mode.

**Figure 6.**
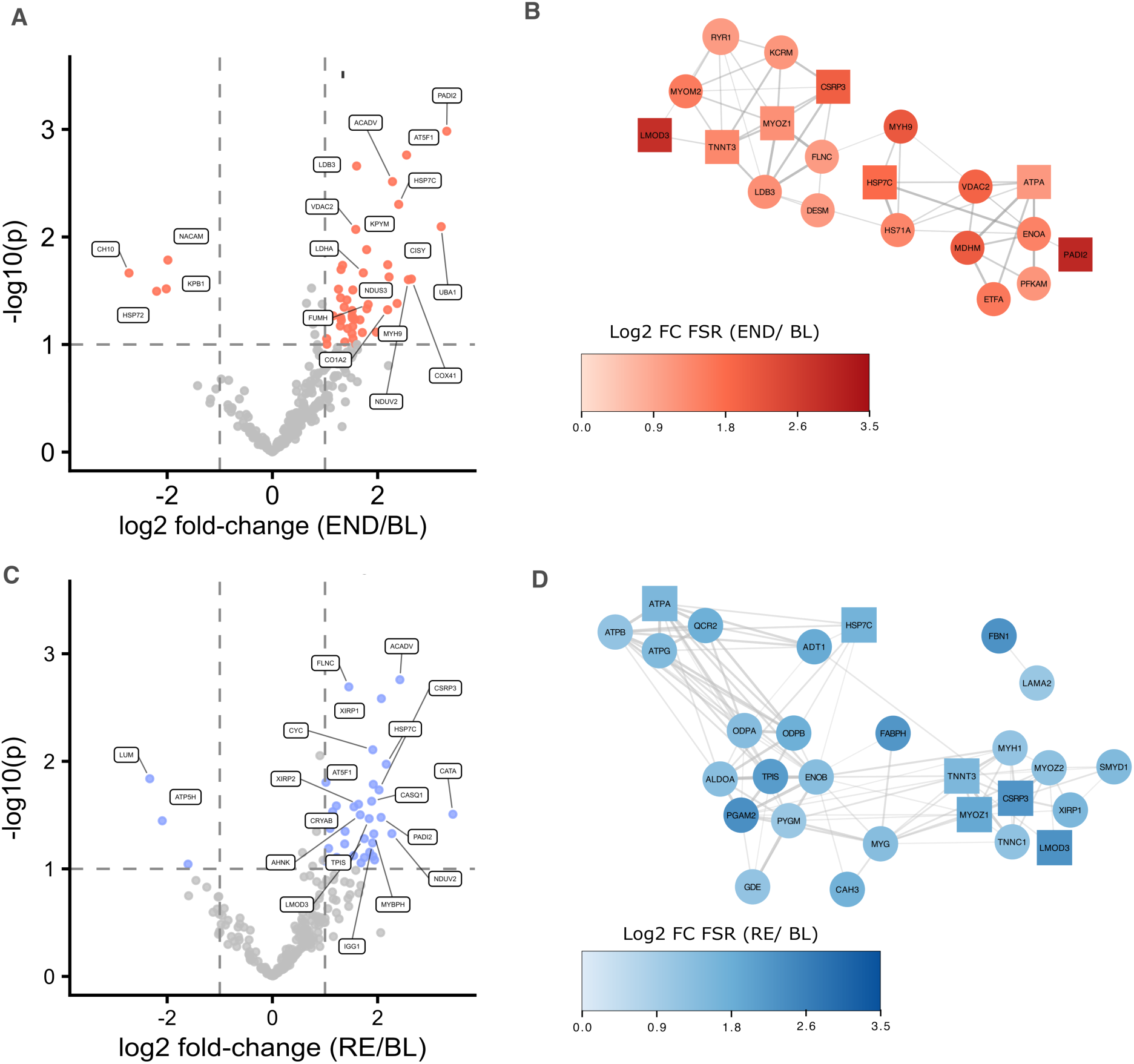
Mode-specific and shared upregulated proteins after 10 weeks of endurance (END) and resistance (RE) training. (A, C) Volcano plots showing protein-specific changes in fractional synthesis rate (FSR; log₂ fold-change vs baseline) following END (A) and RE (C). Significantly regulated proteins (p < 0.1 and log₂FC > 1) are coloured red (END) or blue (RE); grey points indicate non-significant proteins. Labelled proteins represent the top 20 significantly changed FSRs (log₂FC and –log₁₀ p). Dashed lines mark significance thresholds. (B, D) STRING protein–protein interaction networks of upregulated proteins after END (B) and RE (D). Node colour intensity reflects log₂ fold-change in FSR relative to baseline. Circles represent proteins uniquely upregulated in each mode, while squares indicate proteins common to both END and RE. Networks illustrate mode-specific clustering of cytoskeletal, metabolic, and chaperone proteins, with limited overlap in the specific subsets remodelled by each training mode.

The first week of training elicited classical divergent programmes of protein synthesis in skeletal muscle. Protein synthetic responses to early END were dominated by a large network of mitochondrial and metabolic proteins (Figure 5A). A distinct subset of contractile and structural proteins (PDLIM3, LDB3, MYOZ2, TNNT3, SMYD1 and several collagen proteins) increased specifically in END (Figure 5A-B). These data indicate that in untrained muscle, although both RE and END exercise increased mixed-sarcoplasmic and myofibrillar protein synthesis rates, the different training modes selectively increased the synthesis rates of distinct protein subsets within these subcellular fractions.

In contrast, RE induced robust increases in the synthesis of contractile and structural proteins, including MYBPH, MYOZ1, XIRP1 and 2, FLNC, DESM, and DMD (FigureS 5C-D). Regulatory proteins were also affected, such as CSRP3 (a positive regulator of myogenesis) and AHNAK (a neuroblast differentiation-associated protein), suggesting activation of developmental signalling pathways. In addition, RE increased synthesis of key metabolic enzymes involved in rapid ATP resynthesis (KCRM, PYGM, G6PI) alongside mitochondrial and redox enzymes (MDHA, SDHA, HADHA, CATA) (Figure 5D)

Six proteins (ACADV, ATP5F1, CASQ1, CRYAB, CYC, and NDUV2) were increased in synthesis rate in the response to both END and RE across the first week of training (Figure 5B and 5D).

During later adaptation to END, the number of metabolic enzymes with elevated synthesis was reduced compared with Week 1, consistent with a decreased perturbation of metabolic networks in this trained state. However, END maintained a specific subset of Z-disc proteins with increased fractional synthesis rate, including LDB3, FLNC and DESM, suggesting continued structural remodelling of the Z-disc despite no global increase in mixed myofibrillar protein synthesis (Figure 6A-B).

Later RE adaptation was more clearly characterised by increased synthesis of glycolytic enzymes (GDE, PGAM2, PYGM, ALDOA, PDHB, TPIS, ENOB, CAH3), alongside sustained upregulation of structural proteins including fast-twitch myosin heavy chain (MYH1). In addition, a subnetwork of mitochondrial proteins, including ATP synthase subunits (ATPA, ATPB, ATPG), showed increased synthesis during Later RE (Figure 6C-D).

A small group of proteins showed consistently increased synthesis rates across both exercise modes and time points. Three proteins, heat shock cognate 71 kDa protein (HSPA8), leiomodin-3 (LMOD3), and protein-arginine deiminase type-2 (PADI2), were increased across both Early and Later adaptation to END and RE, identifying them as a core conserved exercise response (Figure 7A).

**Figure 7.**
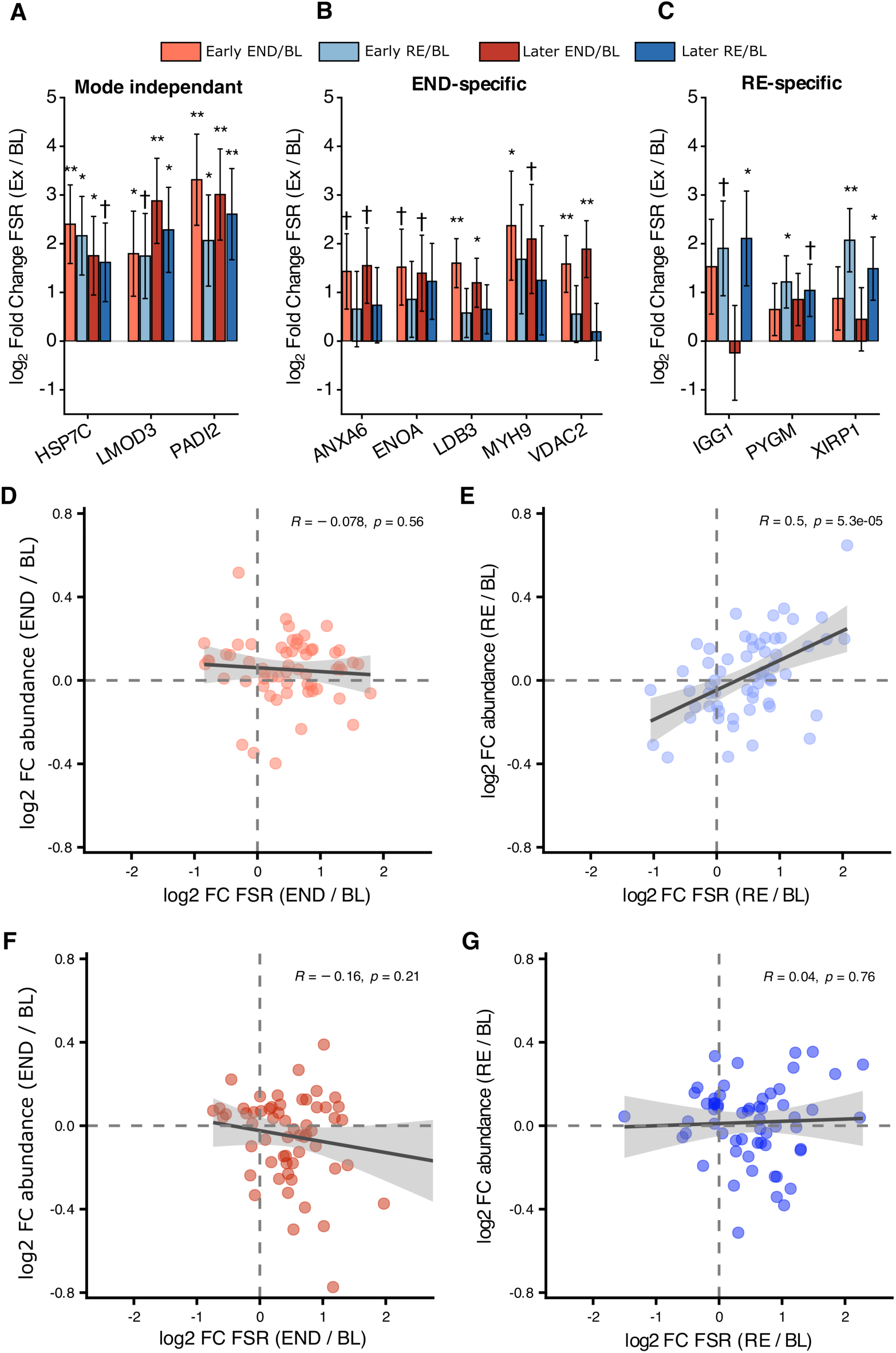
Mode-dependant and independent synthesis responses contribute to muscle adaptation. (A–C) Grouped bar plots show log₂ fold-change in fractional synthesis rate (FSR) relative to baseline (Ex/BL) for selected proteins: (A) mode-independent (HSP7C, LMOD3, PADI2), (B) END-specific (ANXA6, ENOA, HSPA8, LDB3, MYH9, VDAC2), and (C) RE-specific (IGG1, PYGM, XIRP1). Four bars are displayed per protein: Early AE/BL (light red), Early RT/BL (light blue), Later AE/BL (dark red), and Later RT/BL (dark blue). Bars are mean ± SE; horizontal dashed line denotes no change (0 on the log₂ scale). Symbols indicate within-contrast differences from baseline: † p < 0.10, * p < 0.05, ** p < 0.01. (D–G) Protein-level relationships between FSR and abundance (both log₂ fold-change vs baseline) of proteins annotated to the GO Cellular component ‘Myofibril’ with least-squares fit and 95% CI shading. (D, F) END (red) and (E, G) RE (blue), shown for Early (Week 1; D, E) and Later (Week 10; F, G) adaptation. Each point is one protein; dashed lines mark zero on each axis. Pearson’s R and P values are printed in the panel.

Mode-specific conserved changes were also identified. Six proteins (ANXA6, ENOA, HSPA8, LDB3, MYH9, and VDAC2) were consistently upregulated across Early and Later END but not significantly changed in FSR in RE muscle, indicating a mode-specific regulation of their synthesis rate (Figure 7). Conversely, IGG1, PYGM, and XIRP1 showed increased synthesis exclusively in RE muscle across both the Early and Later phases, highlighting RE-specific changes in individual protein synthetic responses (Figure 7). Complete protein-level differential analysis of protein-specific FSR is provided in Supplementary Table 2.

Integration of changes in protein-specific synthesis rates with changes in abundance enables quantitative links between synthesis and abundance during muscle adaptation. Whilst increases in the synthesis rates of individual myofibrillar proteins were identified as a core concept of adaptation to END and RE, correlational analysis revealed changes in synthesis rate did not similarly result in changes in protein abundance. In our analysis, changes in synthesis rate and the abundance of individual myofibrillar proteins were significantly (P < 0.001) positively correlated (R = 0.5) in RE muscle during the early adaptation period, suggesting that RE-induced increases in synthesis contributed directly to quantitative proteome remodelling. Whereas, in END muscle, no such correlation was observed (P > 0.2), implying that increased synthesis primarily contributed to an enhanced turnover rate of muscle proteins rather than net accumulation of myofibrillar proteins. Therefore, the regulation of protein degradation and thus enhanced accumulation or turnover of myofibrillar proteins is a key contributor to training-mode specific adaptation in skeletal muscle

## Discussion

We applied dynamic proteomic profiling to provide the first protein-by-protein characterisation of muscle protein synthesis during END and RE training in humans. Previous studies have largely focused on acute (≤ 24 h) responses to single exercise bouts and have relied on mixed- or fractional (myofibrillar or sarcoplasmic)-specific protein measures, which are unable to uncover the complexity of protein-specific turnover underlying skeletal muscle adaptation. We resolved synthesis rates for 247 individual proteins across the first week (Early) and tenth week (Later) of training. Our data confirmed established exercise mode-specific trends, such as greater myofibrillar protein synthesis with RE and greater sarcoplasmic/mitochondrial synthesis with END. Crucially, our data revealed that distinct protein subsets within the same subcellular location are selectively regulated in an exercise mode-specific manner. For example, RE specifically increased synthesis of contractile and structural proteins, such as MYBPH, MYOZ1, and XIRP1 and XIRP2, whereas END preferentially increased synthesis of distinct Z-disc proteins, including PDLIM3, LDB3, and MYOZ2. Integration of synthesis and abundance data further indicated that RE-induced changes in synthesis contributed to quantitative protein accumulation, whereas END-induced changes in synthesis seemed to reflect enhanced protein turnover. These findings move beyond traditional mixed- and fraction-based analyses to highlight the complexity of exercise-specific proteome adaptation and point to the need for future work on the mechanisms of translational control and, possibly, targeted degradation that shape protein-specific remodelling in response to exercise training.

Consistent with the paradigm that RE stimulates myofibrillar protein synthesis to a greater extent than END in the hours following a single training bout (Wilkinson et al., 2008, Zhu et al., 2025), we observed increased mixed-myofibrillar FSR during Early RE and END, with a greater magnitude of response to RE when measured across 5-days consisting of multiple training bouts (Figure 2). An increased rate of average myofibrillar protein synthesis is a key feature distinguishing RE and END adaptation during both acute (hours) and chronic (days-weeks) adaptation. Interestingly, our dynamic proteomic analyses revealed little concordance between myofibrillar protein synthesis responses between training modes during early adaptation (Figure 4C). Our data indicate that RE preferentially stimulated the synthesis of distinct proteins, such as MYBPH, MYOZ1, XIRP1/2, FLNC, DES, and DMD, whereas END selectively remodelled Z-disc–associated proteins, including PDLIM3, LDB3, MYOZ2, TNNT3, and SMYD1 (Figure 5). Dynamic proteomics is a novel approach offering unique insight into human exercise research, with only a handful of studies published to date, each examining a single training modality in isolation (Shankaran et al., 2016, Camera et al., 2017, Srisawat et al., 2023, Nishimura et al., 2025, Murphy et al., 2018). RE studies have shown increased synthesis of desmin and fast-twitch myosin isoforms (Camera et al., 2017), while sprint interval training elevated synthesis of glycolytic enzymes and cytoskeletal proteins (Shankaran et al., 2016, Srisawat et al., 2023, Nishimura et al., 2025). These mode-specific findings demonstrate that protein synthesis is regulated at the level of individual proteins, providing confidence that direct mode comparisons reveal distinct dynamic signatures. Here, we provide the first direct comparison of proteome-wide synthesis responses to END and RE within the same participant. Although both training modes increased the average synthesis rate of myofibrillar proteins, with a greater magnitude in RE, individual proteins within this fraction were regulated in a mode-dependent manner. This result demonstrates that even within the myofibrillar fraction, averaging across constituent proteins obscures some of the heterogeneous individual protein-level adaptive responses that may contribute to the unique phenotypic outcomes following training. Thus, our data indicate a greater magnitude of myofibrillar protein synthetic response in RE, as well as mode-dependent alterations in translational specificity through the selective synthesis of distinct subsets of myofibrillar proteins, which are associated with, and may be causal in, divergent phenotypic adaptations.

The Z-disc serves as the structural anchor of the sarcomere, integrating mechanical forces across the myofibril and acting as a scaffolding platform for mechanosensitive signalling complexes, making the Z-disc a site of both damage-induced repair and load-driven functional remodelling. Mechanical loading of muscle unaccustomed to exercise, particularly heavy RE, can induce sarcomeric disruption concentrated at the Z-disc (Gibala et al., 1995, Gibala et al., 2000). Importantly, Z-disc disruption is not confined to the immediate post-exercise period, with greater structural disorganisation observed at days 2–3 and 7–8 post-exercise compared with 1 hour post-exercise (Yu et al., 2004). These data indicate that Z-disc remodelling is a potentially longer-term process that influences adaptation beyond the initial bout or response to damage. Our data suggest that Z-disc remodelling is a core feature of adaptation to both END and RE, rather than a response exclusive to what has classically been described as muscle damage. We observed increased synthesis of the α-crystallin B chain (CRYAB) during early adaptation to both RE and END (Figures 5B and 5D), but this was not sustained by either training mode during later adaptation, indicating that enhanced CRYAB synthesis is a conserved response to the onset of novel contractile loading. CRYAB is a small heat shock protein that binds desmin and actin at the Z-disc and maintains myofibrillar proteostasis by protecting against unfolding and aggregation of myosin heavy chains (Melkani et al., 2006). In both animal and human studies, phosphorylation and re-localisation of CRYAB have become widely used markers for structural strain, and sustained increases in CRYAB synthesis have been identified following resistance training (Stead et al., 2025a). Increased CRYAB synthesis is a shared feature of the early adaptive response to both RE and END, suggesting that novel contractile loading, regardless of modality, alters proteostasis. Furthermore, CRYAB may function as a broad regulator of exercise-induced proteostatic rather than a marker of overt muscle damage. Taken together, these findings reposition CRYAB from a classical indicator of exercise-induced muscle damage to a conserved regulator of proteostatic remodelling with unaccustomed exercise training.

In contrast to common responses to RE and END, our data reveal clear mode-specific regulation of individual Z-disc proteins that distinguish RE and END adaptations. Although exercise responses are often framed as structural for RE and metabolic for END, our results show that cytoskeletal remodelling is a broader, mode-dependent feature engaged by both modalities. In RE, we observed a sustained elevation in the synthesis of Xin-actin-binding repeat protein 1 (XIRP1) during both early and later adaptation. Xin is a striated-muscle-specific protein localised to remodelling of Z-discs and myotendinous junctions following high-strain or eccentrically-biased loading (Otten et al., 2012), and it has been widely used as a marker of muscle damage (Nilsson et al., 2013). Our longitudinal data extend this view, showing that XIRP1 abundance increased early in training, consistent with remodelling and repair processes; however, by week 10, there was no change in abundance compared with pre-exercise levels. The protein synthesis rate of XIRP1 remained increased specifically in RE muscle, suggesting that RE-induced increases in XIRP1 turnover are required after early accumulation, highlighting enhanced XIRP1 turnover as a unique and sustained feature of RE-specific cytoskeletal adaptation.

In contrast, END exhibited a selective and sustained increase in LDB3 synthesis, despite no detectable change in abundance, indicating enhanced turnover rather than net accretion. Given the role of LDB3 in anchoring α-actinin and organising Z-disc–linked mechanosensitive complexes (Hoshijima, 2006), increased turnover may reflect ongoing renewal or fine-tuning of Z-disc scaffolding under prolonged cyclic contractions. Whether this increased turnover represents increased quality control, isoform-specific regulation, or modulation of mechanotransduction pathways remains unresolved. Nevertheless, the specificity and sustained nature of LDB3 turnover identify it as an END-associated structural response, mechanistically distinct from the XIRP1 response observed in RE. Collectively, these exercise mode-specific findings demonstrate that Z-disc adaptation is not uniform and that individual proteins within the same structural compartment can exhibit divergent synthesis profiles across exercise training modes. More direct mechanistic work is needed to establish whether these turnover patterns causally contribute to adaptation.

Dynamic proteome profiling provides coordinated measurements of both protein abundance and synthesis rates on a protein-by-protein basis, enabling direct assessment of how changes in synthesis translate into proteome remodelling. When integrating these datasets, we observed no correlation between changes in protein synthesis and abundance in END muscle, implying that increased synthesis primarily contributed to enhanced turnover rather than net protein accretion (Figure 7). In contrast, during early adaptation to RE, increases in fractional synthesis rates of individual myofibrillar proteins were significantly and positively associated with changes in abundance (R = 0.5, P < 0.001) (Figure 7). This alignment indicates that elevated synthesis contributed directly to quantitative remodelling, where increased production of structural proteins results in net accretion of the contractile apparatus. Previous dynamic proteome profiling studies have reported similar patterns, where exercise increases the turnover rate of some proteins during endurance training (Srisawat et al., 2023) and resistance training (Camera et al., 2017). Such increases in turnover (i.e., equivalent gains in synthesis and degradation) improve muscle proteostasis by enhancing protein renewal and preventing the accumulation of damaged or dysfunctional proteins (Goldberg, 2003). However, we observed that the coupling between synthesis and abundance appears to be exercise mode-dependent: RE demonstrated a quantitative link in which elevated synthesis drives net accumulation and muscle growth, whereas END did not. This mode-dependent relationship between changes in synthesis and proteome adaptation directly explains how similar increases in myofibrillar protein synthesis between exercise modes could produce divergent structural and functional phenotypes. Critically, these findings reveal that synthesis rate alone is insufficient to predict the nature or direction of proteome remodelling, at least with END, and that understanding exercise-specific adaptations requires integrating both synthetic and degradative processes.

While dynamic proteomics remains rare in exercise and muscle physiology research, protein abundance profiling is becoming routinely applied to training adaptation, providing a robust literature base against which to compare and validate our findings. After 10 weeks of endurance training, we observed marked increases in proteins associated with mitochondrial function, particularly those localised to the inner mitochondrial membrane and electron transport chain. These observations are consistent with proteomic analyses identifying mitochondrial pathways as the most prominently upregulated modules following endurance exercise (Pataky et al., 2025). Our unilateral END protocol closely mirrored earlier work identifying electron transport chain complex IV as the most robustly upregulated component within skeletal muscle following training (Hostrup, 2021), with significant enrichment of complex IV proteins after 10 weeks (Figure 3). In the resistance-trained limb, early adaptation revealed increased abundance of ribosomal protein networks after one week of RE, supporting recent evidence that ribosomal proteome remodelling (Jessen et al., 2024) and expansion of translational capacity (Stead et al., 2025a) are central components of early-phase adaptation to resistance training. The concordance between our abundance profiles and established mode-specific signatures from independent cohorts qualitatively validates the capacity of our within-subject unilateral design to isolate exercise mode-specific training responses.

Several methodological considerations should be acknowledged when interpreting these findings. The application of false discovery rate (FDR) correction to quantitative proteomics data presents a statistical challenge that warrants careful consideration. FDR thresholds, most commonly applied using the Benjamini–Hochberg procedure, were originally developed for high-dimensional transcriptomic analyses where tens of thousands of hypotheses are tested simultaneously. In that setting, correction is essential, as even a nominal α of 0.05 yields hundreds of expected false positives across the feature space. The statistical landscape of a dynamic proteomics dataset is fundamentally different, requiring stringent longitudinal filtering in the present study, which limited the proteins quantified at every time point across all participants, yielding a substantially smaller analytical set. Under these conditions, global FDR procedures can be disproportionately conservative, as has been formally demonstrated in simulation studies showing that standard multiple testing corrections commonly applied to proteomics data may fail to detect genuine biological effects when the number of features is moderate, and effect sizes are modest (Pascovici et al., 2016), which is typical in human exercise physiology.

Our findings demonstrate that skeletal muscle adapts to RE and END through distinct regulation of proteome dynamics. Early adaptation to RE was characterised by a greater magnitude of myofibrillar protein synthesis, accompanied by the accumulation of ribosomal and contractile proteins, consistent with sustained upregulation of translational capacity and structural remodelling. In contrast, END induced a pattern of remodelling, marked by selective turnover of structural proteins alongside temporal restructuring of mitochondrial sub-compartments, with Early downregulation of matrix proteins preceding Later increases in inner membrane complexes. Importantly, by integrating protein-specific synthesis rates with abundance changes, we showed that RE-induced increases in synthesis contribute directly to protein accumulation. In contrast, in END, synthesis appears more tightly linked to enhanced proteome turnover, particularly for myofibrillar or Z-disc proteins, which exhibited a mode-specific and sustained increase in synthesis rate (e.g., LDB3). Beyond confirming classical fraction-level differences, our data show that subsets of proteins within the same structural class (e.g., Z-disc proteins XIRP1 and LDB3) can be selectively synthesised in a mode-specific manner, underscoring the importance of protein-level resolution for understanding exercise adaptation.

## Availability of data

The mass spectrometry proteomics data have been deposited to the ProteomeXchange Consortium via the PRIDE partner repository with the dataset identifier PXD076386 and [XXXX] for the myofibrillar and soluble fraction, respectively.

## Author contributions

SM Phillips and JG Burniston conceived and designed the research; CA Stead, ACQ Thomas, AC D’Souza, J McKendry, C Lim, JC McLeod, and C McGlory performed the research and acquired the data. CA Stead and ACQ Thomas analysed and interpreted the data. All authors were involved in drafting and revising the manuscript.

## References

Bishop, D. J., Botella, J., Genders, A. J., Lee, M. J., Saner, N. J., Kuang, J., Yan, X. & Granata, C. 2019. High-Intensity Exercise and Mitochondrial Biogenesis: Current Controversies and Future Research Directions. Physiology (Bethesda), 34, 56–70.

Buckley, J. P. & Borg, G. A. 2011. Borg’s scales in strength training; from theory to practice in young and older adults. Appl Physiol Nutr Metab, 36, 682–92.

Burniston, J. G. 2019. Investigating Muscle Protein Turnover on a Protein-by-Protein Basis Using Dynamic Proteome Profiling. In: Burniston, J. G. & Chen, Y.-W. (eds.) Omics Approaches to Understanding Muscle Biology. New York, NY: Springer US.

Burniston, J. G., Connolly, J., Kainulainen, H., Britton, S. L. & Koch, L. G. 2014. Label-free profiling of skeletal muscle using high-definition mass spectrometry. Proteomics, 14, 2339–44.

Camera, D. M., Burniston, J. G., Pogson, M. A., Smiles, W. J. & Hawley, J. A. 2017. Dynamic proteome profiling of individual proteins in human skeletal muscle after a high-fat diet and resistance exercise. Faseb j, 31, 5478–5494.

Damas, F., Libardi, C. A. & Ugrinowitsch, C. 2018. The development of skeletal muscle hypertrophy through resistance training: the role of muscle damage and muscle protein synthesis. Eur J Appl Physiol, 118, 485–500.

Damas, F., Phillips, S. M., Lixandrão, M. E., Vechin, F. C., Libardi, C. A., Roschel, H., Tricoli, V. & Ugrinowitsch, C. 2016. Early resistance training-induced increases in muscle cross-sectional area are concomitant with edema-induced muscle swelling. Eur J Appl Physiol, 116, 49–56.

Egan, B. & Zierath, Juleen R. 2013. Exercise Metabolism and the Molecular Regulation of Skeletal Muscle Adaptation. Cell Metabolism, 17, 162–184.

Emanuelsson, E. B., Arif, M., Reitzner, S. M., Perez, S., Lindholm, M. E., Mardinoglu, A., Daub, C., Sundberg, C. J. & Chapman, M. A. 2024. Remodeling of the human skeletal muscle proteome found after long-term endurance training but not after strength training. iScience, 27.

Gibala, M. J., Interisano, S. A., Tarnopolsky, M. A., Roy, B. D., Macdonald, J. R., Yarasheski, K. E. & Macdougall, J. D. 2000. Myofibrillar disruption following acute concentric and eccentric resistance exercise in strength-trained men. Can J Physiol Pharmacol, 78, 656–61.

Gibala, M. J., Macdougall, J. D., Tarnopolsky, M. A., Stauber, W. T. & Elorriaga, A. 1995. Changes in human skeletal muscle ultrastructure and force production after acute resistance exercise. J Appl Physiol (1985), 78, 702–8.

Goldberg, A. L. 2003. Protein degradation and protection against misfolded or damaged proteins. Nature, 426, 895–899.

Goreham, C., Green, H. J., Ball-Burnett, M. & Ranney, D. 1999. High-resistance training and muscle metabolism during prolonged exercise. Am J Physiol, 276, E489–96.

Hesketh, S. J., Stansfield, B. N., Stead, C. A. & Burniston, J. G. 2021. The application of proteomics in muscle exercise physiology. Expert Rev Proteomics, 17, 813–825.

Hesketh, S. J., Sutherland, H., Lisboa, P. J., Jarvis, J. C. & Burniston, J. G. 2020. Adaptation of rat fast-twitch muscle to endurance activity is underpinned by changes to protein degradation as well as protein synthesis. Faseb j, 34, 10398–10417.

Holmes, W. E., Angel, T. E., Li, K. W. & Hellerstein, M. K. 2015. Dynamic Proteomics: In Vivo Proteome-Wide Measurement of Protein Kinetics Using Metabolic Labeling. Methods Enzymol, 561, 219–76.

Hoshijima, M. 2006. Mechanical stress-strain sensors embedded in cardiac cytoskeleton: Z disk, titin, and associated structures. Am J Physiol Heart Circ Physiol, 290, H1313–25.

Hostrup, M. 2021. High-Intensity Interval Training Remodels the Proteome and Acetylome of Human Skeletal Muscle 2 Hostrup M 1+*, Lemminger Ak 1+, Stocks B 2+, Gonzalez-Franquesa A. Cold Spring Harbor Laboratory.

Husek, P. 1991. Amino acid derivatization and analysis in five minutes. Febs Lett, 280, 354–6.

Ilchenko, S., Haddad, A., Sadana, P., Recchia, F. A., Sadygov, R. G. & Kasumov, T. 2019. Calculation of the Protein Turnover Rate Using the Number of Incorporated 2h Atoms and Proteomics Analysis of a Single Labeled Sample. Analytical Chemistry, 91, 14340–14351.

Jessen, S., Quesada, Júlia P., Di Credico, A., Moreno-Justicia, R., Wilson, R., Jacobson, G., Bangsbo, J., Deshmukh, Atul S. & Hostrup, M. 2024. Beta2-Adrenergic Stimulation Induces Resistance Training-Like Adaptations in Human Skeletal Muscle: Potential Role of KLHL41. Scandinavian Journal of Medicine & Science in Sports, 34, e14736.

Jorgenson, K. W., Phillips, S. M. & Hornberger, T. A. 2020. Identifying the Structural Adaptations that Drive the Mechanical Load-Induced Growth of Skeletal Muscle: A Scoping Review. Cells, 9.

Mcglory, C., White, A., Treins, C., Drust, B., Close, G. L., Maclaren, D. P., Campbell, I. T., Philp, A., Schenk, S., Morton, J. P. & Hamilton, D. L. 2014. Application of the [gamma-32p] Atp kinase assay to study anabolic signaling in human skeletal muscle. J Appl Physiol (1985), 116, 504–13.

Mckendry, J., Lowisz, C. V., Nanthakumar, A., Macdonald, M., Lim, C., Currier, B. S. & Phillips, S. M. 2024. The effects of whey, pea, and collagen protein supplementation beyond the recommended dietary allowance on integrated myofibrillar protein synthetic rates in older males: a randomized controlled trial. Am J Clin Nutr.

Mckendry, J., Shad, B. J., Smeuninx, B., Oikawa, S. Y., Wallis, G., Greig, C., Phillips, S. M. & Breen, L. 2019. Comparable Rates of Integrated Myofibrillar Protein Synthesis Between Endurance-Trained Master Athletes and Untrained Older Individuals. Front Physiol, 10, 1084.

Mcleod, J. C., Currier, B. S., Lowisz, C. V. & Phillips, S. M. 2024. The influence of resistance exercise training prescription variables on skeletal muscle mass, strength, and physical function in healthy adults: An umbrella review. Journal of Sport and Health Science, 13, 47–60.

Melkani, G. C., Cammarato, A. & Bernstein, S. I. 2006. alphab-crystallin maintains skeletal muscle myosin enzymatic activity and prevents its aggregation under heat-shock stress. J Mol Biol, 358, 635–45.

Mølmen, K. S., Almquist, N. W. & Skattebo, Ø. 2025. Effects of Exercise Training on Mitochondrial and Capillary Growth in Human Skeletal Muscle: A Systematic Review and Meta-Regression. Sports Med, 55, 115–144.

Murphy, C. H., Shankaran, M., Churchward-Venne, T. A., Mitchell, C. J., Kolar, N. M., Burke, L. M., Hawley, J. A., Kassis, A., Karagounis, L. G., Li, K., King, C., Hellerstein, M. & Phillips, S. M. 2018. Effect of resistance training and protein intake pattern on myofibrillar protein synthesis and proteome kinetics in older men in energy restriction. John Wiley & Sons, Ltd.

Nilsson, M. I., Nissar, A. A., Al-Sajee, D., Tarnopolsky, M. A., Parise, G., Lach, B., Fürst, D. O., Van Der Ven, P. F. M., Kley, R. A. & Hawke, T. J. 2013. Xin Is a Marker of Skeletal Muscle Damage Severity in Myopathies. The American Journal of Pathology, 183, 1703–1709.

Nishimura, Y., Bittel, A. J., Stead, C. A., Chen, Y. W. & Burniston, J. G. 2023. Facioscapulohumeral Muscular Dystrophy is Associated With Altered Myoblast Proteome Dynamics. Mol Cell Proteomics, 22, 100605.

Nishimura, Y., Langan-Evans, C., Taylor, H. L., Foo, W. L., Morton, J. P., Shepherd, S., Strauss, J. A., Burniston, J. G. & Areta, J. L. 2025. Endocrine, Metabolic, and Skeletal Muscle Proteomic Responses During Energy Deficit With Concomitant Aerobic Exercise in Humans. Faseb j, 39, e71163.

Pascovici, D., Handler, D. C., Wu, J. X. & Haynes, P. A. 2016. Multiple testing corrections in quantitative proteomics: A useful but blunt tool. Proteomics, 16, 2448–53.

Pataky, M. W., Heppelmann, C. J., Sevits, K. J., Asokan, A. K., Kumar, A. P., Klaus, K. A., Dasari, S., Kunz, H. E., Strub, M. D., Robinson, M. M., Coon, J. J., Lanza, I. R., Adams, C. M. & Nair, K. S. 2025. Aerobic and resistance exercise-regulated phosphoproteome and acetylproteome modifications in human skeletal muscle. Nature Communications, 16, 5700.

Roberts, M. D., Mccarthy, J. J., Hornberger, T. A., Phillips, S. M., Mackey, A. L., Nader, G. A., Boppart, M. D., Kavazis, A. N., Reidy, P. T., Ogasawara, R., Libardi, C. A., Ugrinowitsch, C., Booth, F. W. & Esser, K. A. 2023. Mechanisms of mechanical overload-induced skeletal muscle hypertrophy: current understanding and future directions. Physiol Rev, 103, 2679–2757.

Sadygov, R. G. 2020. Partial Isotope Profiles Are Sufficient for Protein Turnover Analysis Using Closed-Form Equations of Mass Isotopomer Dynamics. Anal Chem, 92, 14747–14753.

Shankaran, M., King, C. L., Angel, T. E., Holmes, W. E., Li, K. W., Colangelo, M., Price, J. C., Turner, S. M., Bell, C., Hamilton, K. L., Miller, B. F. & Hellerstein, M. K. 2016. Circulating protein synthesis rates reveal skeletal muscle proteome dynamics. J Clin Invest, 126, 288–302.

Shannon, P., Markiel, A., Ozier, O., Baliga, N. S., Wang, J. T., Ramage, D., Amin, N., Schwikowski, B. & Ideker, T. 2003. Cytoscape: a software environment for integrated models of biomolecular interaction networks. Genome Res, 13, 2498–504.

Srisawat, K., Stead, C. A., Hesketh, K., Pogson, M., Strauss, J. A., Cocks, M., Siekmann, I., Phillips, S. M., Lisboa, P. J., Shepherd, S. & Burniston, J. G. 2023. People with obesity exhibit losses in muscle proteostasis that are partly improved by exercise training. Proteomics, e2300395.

Stead, C. A., Hesketh, S. J., Thomas, A. C. Q., Viggars, M. R., Sutherland, H., Jarvis, J. C. & Burniston, J. G. 2025a. Dynamic time course of muscle proteome adaptation to programmed resistance training in rats. Am J Physiol Cell Physiol.

Stead, C. A., Thomas, A., Nishimura, Y., Abbasi, M., Barrett, J. & Burniston, J. G. 2025b. Muscle Proteome Dynamics. Adv Exp Med Biol, 1478, 113–153.

Stokes, T., Tripp, T. R., Murphy, K., Morton, R. W., Oikawa, S. Y., Lam Choi, H., Mcgrath, J., Mcglory, C., Macdonald, M. J. & Phillips, S. M. 2021. Methodological considerations for and validation of the ultrasonographic determination of human skeletal muscle hypertrophy and atrophy. Physiol Rep, 9, e14683.

Storey, J. D. & Tibshirani, R. 2003. Statistical significance for genomewide studies. Proc Natl Acad Sci U S A, 100, 9440–5.

Thomas, A. C. Q., Brown, A., Hatt, A. A., Manta, K., Costa-Parke, A., Kamal, M., Joanisse, S., Mcglory, C., Phillips, S. M., Kumbhare, D. & Parise, G. 2022. Short-term aerobic conditioning prior to resistance training augments muscle hypertrophy and satellite cell content in healthy young men and women. Faseb J, 36, e22500.

Wackerhage, H., Schoenfeld, B. J., Hamilton, D. L., Lehti, M. & Hulmi, J. J. 2019. Stimuli and sensors that initiate skeletal muscle hypertrophy following resistance exercise. J Appl Physiol (1985), 126, 30–43.

Wilkinson, D. J., Franchi, M. V., Brook, M. S., Narici, M. V., Williams, J. P., Mitchell, W. K., Szewczyk, N. J., Greenhaff, P. L., Atherton, P. J. & Smith, K. 2014. A validation of the application of D(2)O stable isotope tracer techniques for monitoring day-to-day changes in muscle protein subfraction synthesis in humans. Am J Physiol Endocrinol Metab, 306, E571–9.

Wilkinson, S. B., Phillips, S. M., Atherton, P. J., Patel, R., Yarasheski, K. E., Tarnopolsky, M. A. & Rennie, M. J. 2008. Differential effects of resistance and endurance exercise in the fed state on signalling molecule phosphorylation and protein synthesis in human muscle. J Physiol, 586, 3701–17.

Wiśniewski, J. R., Zougman, A., Nagaraj, N. & Mann, M. 2009. Universal sample preparation method for proteome analysis. Nat Methods, 6, 359–62.

Yu, G., Wang, L. G., Han, Y. & He, Q. Y. 2012. clusterProfiler: an R package for comparing biological themes among gene clusters. Omics, 16, 284–7.

Yu, J. G., Carlsson, L. & Thornell, L. E. 2004. Evidence for myofibril remodeling as opposed to myofibril damage in human muscles with Doms: an ultrastructural and immunoelectron microscopic study. Histochem Cell Biol, 121, 219–27.

Zhu, W. G., Thomas, A. C. Q., Wilson, G. M., Mcglory, C., Hibbert, J. E., Flynn, C. G., Sayed, R. K. A., Paez, H. G., Meinhold, M., Jorgenson, K. W., You, J. S., Steinert, N. D., Lin, K. H., Macinnis, M. J., Coon, J. J., Phillips, S. M. & Hornberger, T. A. 2025. Identification of a resistance-exercise-specific signalling pathway that drives skeletal muscle growth. Nat Metab.

